# CHASMplus reveals the scope of somatic missense mutations driving human cancers

**DOI:** 10.1101/313296

**Authors:** Collin Tokheim, Rachel Karchin

## Abstract

Large-scale cancer sequencing studies of patient cohorts have statistically implicated many genes driving cancer growth and progression, and their identification has yielded substantial translational impact. However, a remaining challenge is to increase the resolution of driver prediction from the gene level to the mutation level, because mutation-level predictions are more closely aligned with the goal of precision cancer medicine. Here we present CHASMplus, a computational method, that is uniquely capable of identifying driver missense *mutations*, including those specific to a cancer type, as evidenced by significantly superior performance on diverse benchmarks. Applied to 8,657 tumor samples across 32 cancer types in The Cancer Genome Atlas, CHASMplus identifies over 4,000 unique driver missense mutations in 240 genes, supporting a prominent role for rare driver mutations. We show which TCGA cancer types are likely to yield discovery of new driver missense mutations by additional sequencing, which has important implications for public policy.

**Significance:** Missense mutations are the most frequent mutation type in cancers and the most difficult to interpret. While many computational methods have been developed to predict whether genes are cancer drivers or whether missense mutations are generally deleterious or pathogenic, there has not previously been a method to score the oncogenic impact of a missense mutation specifically by cancer type, limiting adoption of computational missense mutation predictors in the clinic. Cancer patients are routinely sequenced with targeted panels of cancer driver genes, but such genes contain a mixture of driver and passenger missense mutations which differ by cancer type. A patient’s therapeutic response to drugs and optimal assignment to a clinical trial depends on both the specific mutation in the gene of interest and cancer type. We present a new machine learning method honed for each TCGA cancer type, and a resource for fast lookup of the cancer-specific driver propensity of every possible missense mutation in the human exome.

## Introduction

Cancer is a disease of the genome where certain somatic mutations drive the neoplastic process of cancer, while the majority of mutations are benign passengers (Tomasetti et al., 2015). Missense mutations are the most common protein-coding mutation found in cancer genomes (Vogelstein et al., 2013). A growing set of missense mutations has been recognized as clinically actionable (Hyman et al., 2017b) and more putatively actionable mutations will likely be verified in the future (Bailey et al., 2018). However, due to the large number of somatic mutations identified in DNA sequencing of human tumors, it has been logistically impossible to experimentally validate driver mutations at sufficiently large scale. The task of identifying putative drivers from cancer sequencing studies has therefore depended on statistical models to identify genes with an excess number of mutations over expectation (Davoli et al., 2013; Dees et al., 2012; Lawrence et al., 2013). The rationale is that because driver mutations provide a fitness advantage to cancer cells, they will be observed more often than expected by chance due to their contribution to carcinogenesis and, as such, leave a statistically detectable signal (Kandoth et al., 2013; Lawrence et al., 2014; Martincorena et al., 2017; Parsons et al., 2008; Vogelstein et al., 2013). Based on such gene-level analyses, it has been hypothesized that cancer driver mutations exhibit a long tail phenomenon with few common drivers and many rare drivers(Ding et al., 2010; Garraway and Lander, 2013), suggesting that numerous rare drivers remain to be discovered.

In progressively more hospitals, cancer patients’ tumors are now routinely sequenced with targeted panels of cancer driver genes (*e.g*., MSK-IMPACT, OncoPanel, and many others (Consortium, 2017)). However, even well-established cancer driver genes are thought to contain a mixture of driver and passenger mutations when examined across many patients’ tumors (Vogelstein et al., 2013). This leads to uncertainty on whether an individual mutation is a driver. Attempts have been made to improve the specificity of driver discovery by focusing on smaller intra-genic regions, such as protein domains(Yang et al., 2015), protein-protein interfaces(Engin et al., 2016; Porta-Pardo et al., 2015), and individual codons(Chang et al., 2016). The relevance of shifting from gene-level to mutation-level interpretation is apparent in the American College of Medical Genetics and Genomics (ACMG) guidelines for clinical laboratories (Richards et al., 2015). In the guidelines, which are followed by all U.S. medical genetics laboratories, knowledge of the gene in which a missense mutation is found provides weak evidence for its disease impact, substantially lower than “moderate”, “strong” or “very strong” levels of evidence. Thus, it is recognized that the gene alone is insufficient to ascertain disease impact of a missense mutation. These guidelines were designed for classification of germline missense variants but are consistent with established somatic variant guidelines (Li et al., 2017). Overall, this highlights the critical importance of new methods that can identify putative driver missense mutations and separate them from passenger mutations even within known cancer genes, which could spur effective experimental validation.

While many computational methods have been developed to predict whether a missense mutation is generally deleterious or pathogenic (Adzhubei et al., 2010; Carter et al., 2013; Ioannidis et al., 2016; Jagadeesh et al., 2016; Kircher et al., 2014; Ng and Henikoff, 2001), there has not previously been a method to score the oncogenic impact of a missense mutation specifically by cancer type. Although it is well known that missense mutations have different impacts in different cancer types, currently available computational methods (Adzhubei et al., 2010; Carter et al., 2009; Carter et al., 2013; Cohen et al., 2018; Gonzalez-Perez et al., 2012; Ioannidis et al., 2016; Jagadeesh et al., 2016; Kumar et al., 2016; Mao et al., 2013; Ng and Henikoff, 2001; Reva et al., 2011; Shihab et al., 2013) either do not take it into consideration or fail at the task of distinguishing the differences. A recent systematic study comparing 15 such methods concluded that none of them were yet sufficiently reliable to guide high-cost experimental or clinical follow-through(Martelotto et al., 2014). This indicates an unmet need, given the clinical relevance of mutations is increasingly understood to depend both on the particular mutation and cancer type (Brose et al., 2016; Hyman et al., 2018; Hyman et al., 2017a; Prahallad et al., 2012).

In this work, we present a new machine learning method, CHASMplus, to predict the driver status of missense mutations in a cancer type-specific manner. After careful benchmarking (Result), we applied CHASMplus to 8,657 sequenced tumors from The Cancer Genome Atlas (TCGA) spanning 32 types of cancer, using a statistically rigorous mutational background model to control false discoveries. We explore the emerging role for rare driver missense mutations in cancer and, when possible, relate predictions to supporting functional evidence. We provide an interactive resource for exploring driver missense mutations identified from the TCGA (http://karchinlab.github.io/CHASMplus) and a user-friendly tool (http://chasmplus.readthedocs.io/) to predict whether newly observed mutations from further sequencing are likely cancer drivers. Lastly, we examine the diversity of driver missense mutations across various types of cancer, which leads to a refined understanding of the likely trajectory of driver missense mutation discovery with further sequencing.

## Result

### Overview of CHASMplus

We have developed a new method named CHASMplus that uses machine learning to discriminate somatic missense mutations (referred to hereafter as *missense mutations*) as either cancer drivers or passengers (Figure 1a, Methods). In contrast to our recent analysis of The Cancer Genome Atlas (TCGA) mutations (Bailey et al., 2018), the new method is designed so that predictions can be done in a *cancer type-specific manner* (Figure 1b), as opposed to only considered across multiple cancer types in aggregate (“pan-cancer”). To generate predictions, CHASMplus is trained using somatic mutation calls from TCGA covering 8,657 samples in 32 cancer types (Figure S1, Table S1, Methods). Because there is no gold standard set of driver and passenger missense mutations, we developed a semi-supervised approach to assign class labels to missense mutations. Finally, mutation scores from CHASMplus are weighted by a driver gene score for the respective gene, producing gene-weighted (gwCHASMplus) scores (Methods).

**Figure 1.**
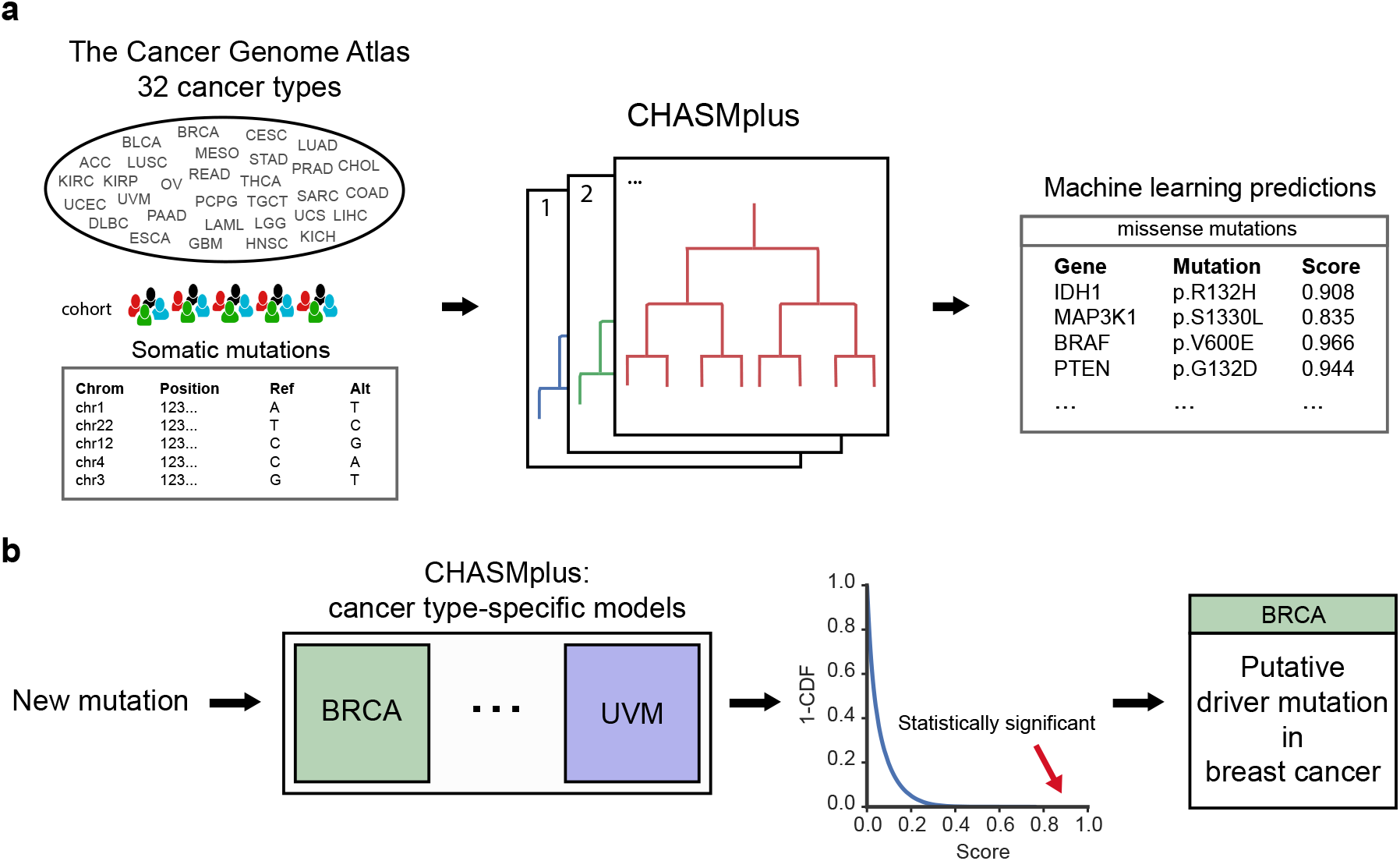
Overview of CHASMplus. **a)** CHASMplus, a machine learning approach, was applied to somatic missense mutations in tumors from 32 different cancer types found in The Cancer Genome Atlas (TCGA). High scores predicted by CHASMplus indicate stronger evidence for a mutation being a cancer driver. **b)** CHASMplus has a model for each cancer type in the TCGA to identify driver missense mutations specific to a cancer type. Putative driver missense mutations are identified at a False Discovery Rate (FDR) threshold of 1 %. In the example, the mutation was statistically significant for the BRCA model but not for the UVM model. Abbreviations: Breast Invasive Carcinoma (BRCA), Uveal Melanoma (UVM) and cumulative distribution function (CDF).

### CHASMplus predicts cancer type-specificity of driver missense mutations

CHASMplus provides a predictive model for each of 32 cancer types sequenced by TCGA. In contrast, most previous methods provide a single impact score for each missense mutation (Adzhubei et al., 2010; Carter et al., 2013; Gonzalez-Perez et al., 2012; Ioannidis et al., 2016; Jagadeesh et al., 2016; Kumar et al., 2016; Ng and Henikoff, 2001; Reva et al., 2011; Shihab et al., 2013), regardless of cancer type. However, two methods (CHASM (Carter et al., 2009) and CanDrA (Mao et al., 2013)) do provide cancer type-specific prediction models, but this capability has not been validated. To illustrate the significant advance in cancer-specific prediction made by CHASMplus, we compared the cancer type-specificity of CHASMplus to CHASM and CanDrA, along with, for reference, two additional methods (ParsSNP (Kumar et al., 2016) and REVEL (Ioannidis et al., 2016)) that are not cancer type-specific.

First, a cancer type-specific model should accurately predict oncogenic effects of missense mutations in an appropriate cell line (Fröhling et al., 2007; Wan et al., 2004). We therefore compared predictions of breast cancer-specific CHASMplus, CHASM, and CanDrA models in known breast cancer driver genes to a previous large-scale validation of 698 missense mutations in MCF10A (breast epithelium) cells that measured cell viability (Ng et al., 2018) (Figure 2a, Methods). We used the area under the Receiver Operating Characteristic curve (auROC) as a performance metric, similar to many prior studies of variant effect prediction (Adzhubei et al., 2010; Ioannidis et al., 2016; Kircher et al., 2014; Kumar et al., 2016; Mao et al., 2013). In general, auROC values range from 0.5 (random prediction performance) to 1.0 (perfect). We found that CHASMplus had substantially higher auROC than compared to CHASM and CanDrA (p<2.2e-16, DeLong test, Table S2). It was also significantly higher than ParsSNP, which is not cancer type-specific, and REVEL, a general-purpose pathogenicity predictor (p<2.2e-16, DeLong test). In fact, CanDrA and CHASM had a lower auROC than ParsSNP, suggesting that these prior methods only captured a limited amount of cancer type-specificity.

**Figure 2.**
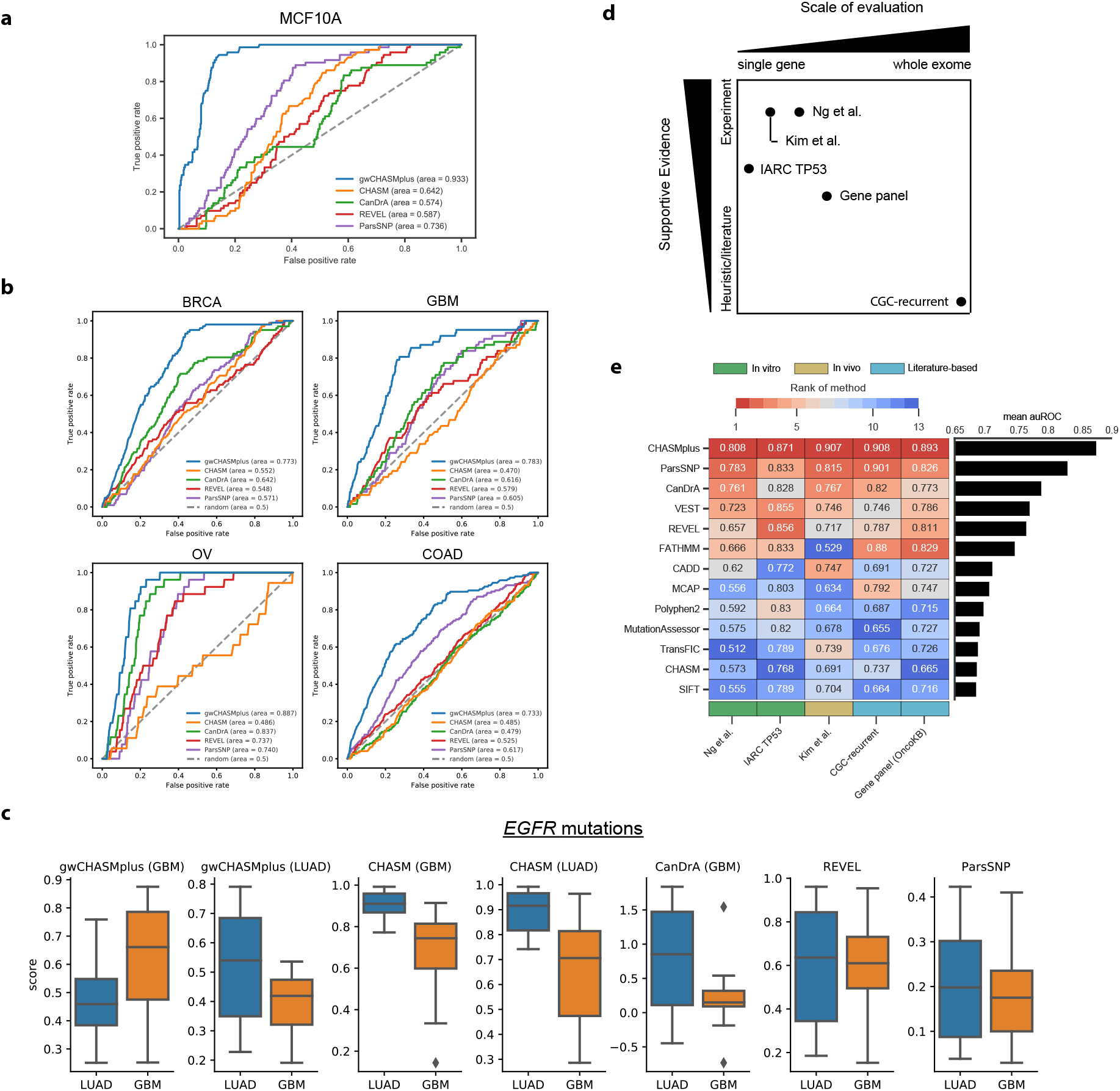
Assessment of pan-cancer and cancer type-specific predictions. **a)** Receiver Operating Characteristic (ROC) curve for identifying activating mutations in MCF10A cells, a breast epithelium line, with breast cancer-specific models. **b)** ROC curve for discerning literature annotated oncogenic mutations from OncoKB for specific cancer types: Breast Invasive Ductal Carcinoma (BRCA), Glioblastoma Multiforme (GBM), High-Grade Serous Ovarian Cancer (OV), and Colon Adenocarcinoma (COAD). **c)** Score distribution for cancer type-specific models (labeled by title) for TCGA missense mutations in *EGFR*, either found in lung adenocarcinoma (LUAD, typically kinase domain mutation) or glioblastoma multiforme (GBM, typically extracellular domain mutations). **d)** Overview of 5 pan-cancer benchmarks by scale of evaluation and type of supportive evidence. **e)** Pan-cancer performance measured by the area under the Receiver Operating Characteristic Curve (auROC) on the 5 mutation-level benchmarks (shown in text). The color scale from red to blue indicates methods ranked from high to low performance. Benchmarks are categorized by *in vitro* (green), *in vivo* (yellow), and literature-based benchmarks (turquoise). The bar graph shows the mean auROC across the benchmarks.

A cancer type-specific model should also be able to distinguish the relevant cancer type among driver mutations in human tumors. We therefore used a literature curated mutation database (OncoKB, (Chakravarty et al., 2017)) to annotate an independent cohort of ~10,000 patients whose tumors were sequenced on a targeted gene panel (MSK-IMPACT, (Zehir et al., 2017)) for oncogenic mutations (Methods). We compared performance on four cancer types (Breast Invasive Ductal Carcinoma [BRCA], Glioblastoma Multiforme [GBM], High-Grade Serous Ovarian Cancer [OV], and Colon Adenocarcinoma [COAD]), as these overlapped cancer types found in the TCGA, and CHASMplus, CHASM and CanDrA had cancer-specific models for these types. CHASMplus had a significantly higher auROC compared to all other methods for each of the cancer types (p<0.05, DeLong test, Figure 2b, Table S2). CanDrA performed better than ParsSNP or REVEL on OV, but generally neither CanDrA nor CHASM showed consistent improvements over ParsSNP and REVEL.

Lastly, we reasoned that distinguishing cancer type-specificity of driver mutations within the same gene would be an even harder task to accomplish. It has been previously documented that lung adenocarcinoma (LUAD) missense mutations in *EGFR* appear predominantly in its kinase domain, while GBM missense mutations appear in its extracellular domain (Brennan et al., 2013; Ji et al., 2006; Paez et al., 2004; Porta-Pardo et al., 2017). We therefore scored TCGA missense mutations in the gene *EGFR* from LUAD patients and from GBM patients with CHASMplus, CanDrA, and CHASM (Figure 2c). The CHASMplus GBM model correctly scores the missense mutations from GBM patients significantly higher than those from LUAD patients (p=0.004, two-sided t-test), and vice versa for the CHASMplus LUAD model (p=0.003, two-sided t-test). In contrast, the CHASM GBM model and the CHASM LUAD model both score the mutations from LUAD patients higher than those from GBM patients (p=1e-5 and 5e-5, respectively, two-sided t-test). CanDrA does not have a LUAD model, but its GBM model scores mutations from LUAD patients higher than those from GBM patients (p=0.0002, twosided t-test), which is significant in the wrong direction. Both REVEL and ParsSNP showed no significant differences in scores between GBM and LUAD (p=0.99 and 0.53, respectively, twosided t-test).

To our knowledge, CHASMplus is the first computational method that can distinguish between driver and passenger missense mutations, specifically by cancer type. Methods published previously have been useful for distinguishing driver mutations, in general, but were limited by the lack of available data for individual cancer types. The poor performance of the 4 other methods tested here, which are among the best in the field, illustrates the substantial improvement by CHASMplus.

### CHASMplus improves pan-cancer identification of driver missense mutations

In contrast to cancer type-specific approaches, there are many excellent existing methods that have been used for pan-cancer analysis. This approach is useful because some cancer driver mutations do occur in many cancer types. The power to detect these mutations, particularly when they occur at low frequency in many cancer types, is increased when many cancer types are aggregated, known as a pan-cancer analysis (Cancer Genome Atlas Research et al., 2013).

We next sought to evaluate whether CHASMplus would also perform well in a pan-cancer analysis, where mutations from all cancer types were modeled together. Because of the greater breadth of relevant methods, we were able to conduct a much broader comparison and a larger number of published benchmarks were available. We compared CHASMplus to 12 methods that span different computational approaches, including those that predict protein functional damage or pathogenicity, and selected meta-predictors (aggregates of multiple methods) -- such as M-CAP (Jagadeesh et al., 2016), REVEL (Ioannidis et al., 2016), and ParsSNP (Kumar et al., 2016) – based on performance in a recent comparative study (Ghosh et al., 2017). We compared these methods on 5 benchmarks, which fall under three broad categories: *in vitro* experiments, a high throughput *in vivo* screen, and curation from published literature. Each of these categories has weaknesses, but, in aggregate, they span multiple scales of evaluation and type of supportive evidence (Figure 2d). For example, several benchmarks were limited to one or a few well-established driver genes, while others were exome-wide but lacked experimental support. A range of benchmarks was critical because missense mutations with the most established experimental support for a driver role tend to be in a few well-understood cancer driver genes. However, limiting benchmarking to these genes would make it difficult to assess the generalizability of a method’s performance to missense mutations in other genes.

Overall, CHASMplus had the highest auROC on each benchmark (Figure 2e), and these differences were statistically significant in all comparisons (p<0.05, DeLong Test, Figure S2b-f, Table S2, Methods). An alternative metric called the area under the Precision-Recall curve yielded similar conclusions as auROC (Figure S2g-i, Methods). These results suggest that CHASMplus improves on previous methods to predict driver missense mutations in a pan-cancer setting, as well as in a cancer type-specific manner.

### CHASMplus identifies putative driver mutations in 32 cancer types

The TCGA genomic landscape across many cancer types has been analyzed with respect to driver genes and, also, with respect to pan-cancer identification of driver mutations. Based on our novel cancer type-specific predictions and improvements in pan-cancer analysis, we hypothesized that new insights could be revealed by CHASMplus. Using cancer type-specific models, we found a wide range of putative driver missense mutations in various cancer types. The lowest was 8 in thymoma, while the highest was 572 in bladder urothelial carcinoma (median of 78, Figure S3a-b, Table S3). In total, 479 unique driver missense mutations were identified by the cancer type-specific analyses but missed by pan-cancer analysis. The pan-cancer analysis identified 3,527 unique missense mutations as putative drivers (Table S3) and had substantial overlap with our earlier pan-cancer analysis (p<2.2e-16, one-sided Fisher’s exact test) (Bailey et al., 2018). Given the substantially different methodology, the overlap suggests the CHASMplus pan-cancer analysis identifies a core set of driver mutations with multiple distinct sources of evidence. Moreover, the pan-cancer analysis also identified 1,369 putative drivers that were missed by the cancer type-specific analysis, supporting our claim that it may be useful to do both.

Interestingly, the cancer type-specific analysis identified a substantial number of putative driver mutations not previously characterized in OncoKB (median overlap 53%). Altogether across the pan-cancer and cancer type-specific analyses, 4,006 unique driver missense mutations were identified by CHASMplus, of which 2,037 were neither found by OncoKB nor our earlier pan-cancer analysis (Bailey et al., 2018), indicating a potentially expanded landscape of putative driver missense mutations of interest for further examination. While it is possible that this could arise from false positives from CHASMplus, our extensive benchmarks (Figure 2) and well-calibrated statistical model (Figure S1f) would suggest this is not the case.

Although the focus of CHASMplus is driver missense mutations, of the 240 genes containing at least one predicted driver missense mutation, 75 genes may be novel. They were not previously included in either the Cancer Gene Census (Forbes et al., 2017) or our previous work with the TCGA PancanAtlas (Bailey et al., 2018) (Table S4). Interestingly, based on gene ontology enrichment (Huang et al., 2008), these genes occurred in pathways related to known hallmarks of cancer (Hanahan and Weinberg, 2011), such as biological processes for DNA damage response, cell cycle, transcriptional regulation, cell-cell adhesion, and cell differentiation (Table S4, Methods). These results suggest that CHASMplus has potential to discriminate driver and passenger mutations in both well-known and putative cancer driver genes, although follow up experiments are required for confirmation.

### CHASMplus identifies both common and rare cancer driver mutations

Previous driver gene studies have suggested that there are few common driver genes and many rare driver genes (long tail hypothesis (Ding et al., 2010; Garraway and Lander, 2013)). However, the overall mutation frequency of a gene does not account for the confounding presence of passenger mutations within a driver gene. Based on our mutation-level analysis, we observed that the spectrum of rare (<1% of cancer samples), intermediate (1-5%), and common (>5%) frequency driver missense mutations varied substantially among cancer types (Figure 3a). For example, uveal melanoma was dominated by common driver missense mutations (88%), while head and neck squamous cell carcinoma (HNSC) was dominated by rare driver missense mutations (63%). However, when we considered all cancer types together (pan-cancer), the overall proportion of rare driver missense mutations considered rare was slightly greater than for common (35.5% and 35.4%, respectively) and 4-fold greater than found by the cancer hotspots method (8%, P<2.2e-16, Fisher’s exact test)(Chang et al., 2016). These results suggest that rare driver missense mutations have a greater role in many cancer types than previously suggested, but that this might not be the case for every cancer type. We observed, after adjusting for sample size, that the average tumor mutation burden for a cancer type positively correlated with the prevalence of rare (but not common) driver missense mutations (R=0.63, P=4.7e-5, likelihood ratio test, Figure 3b).

**Figure 3.**
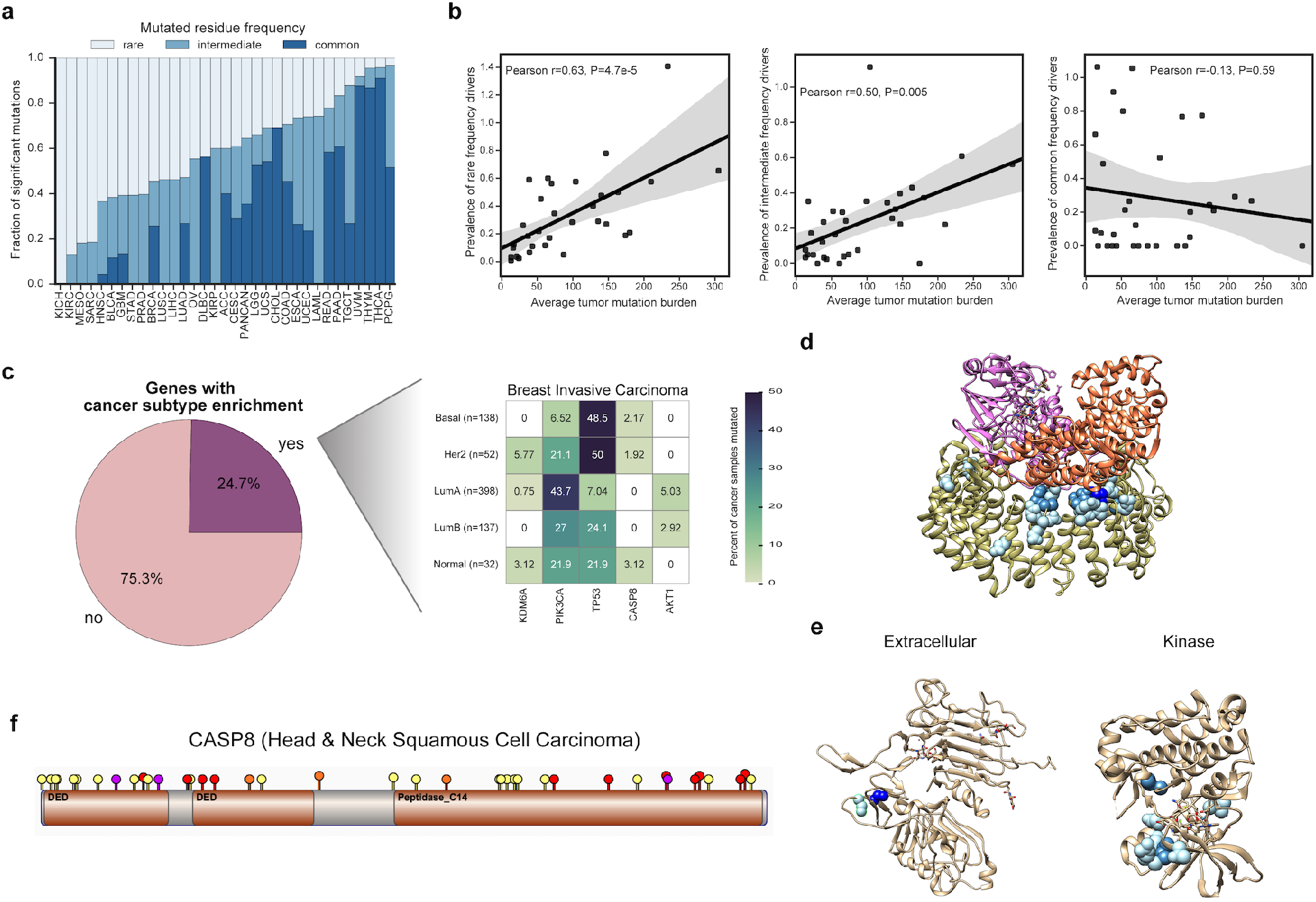
Frequency spectrum of driver missense mutations. **a)** Proportion of the overall frequency of driver missense mutations found to be rare (<1 *%* of samples or singleton mutations), intermediate (1-5%), and common (>5%). Correspondingly shown as light to dark blue. **b)** Correlation between tumor mutation burden and overall driver prevalence (number of driver mutations per tumor) across the frequency spectrum of drivers (rare, intermediate, and common). Shaded area indicates the 95% confidence interval. **c)** Analysis of genes containing predicted driver missense mutations that preferentially occurred in a subtype-specific manner (Chi-squared Test, q<0.1). The pie chart illustrates the percentage of genes for all cancer types with a significant subtype-specific pattern, while the heatmap illustrates significant genes for Breast Invasive Carcinoma. **d)** Structure of the Phosphatase 2A holoenzyme (PDB 2IAE). **e)** Structures of the ERBB2 extracellular domain (left, PDB 2A91) and kinase domain (right, PDB 3PP0). **f)** Lollipop plot of driver missense mutations identified by CHASMplus (yellow), and likely truncating variants (frameshift insertion or deletion: purple, nonsense mutation: red, and splice site mutation: orange) in *CASP8* for Head & Neck squamous cell carcinoma (HNSC). TCGA acronyms for cancer types are listed in methods.

While cancer types appear to have different proportions of rare, intermediate and common driver missense mutations across cancer types, this result could be confounded by differences in subtypes. A common driver mutation in an uncommon subtype, could be perceived, overall, as rare. To test this, we analyzed whether driver missense mutations within a gene showed noticeable enrichment for samples of a particular subtype. In TCGA there are 12 cancer types with available subtype information (Sanchez-Vega et al., 2018). Subtype-specificity partially explained differences in driver mutation frequency spectrum between cancer types. Fifty-five out of 223 genes (24.7%) contained putative driver missense mutations that were enriched in particular cancer subtypes (q-value≤0.1, chi-squared test, Figure 3c, Table S5, Figure S4). Several genes showed strong subtype specificity, consistent with prior literature, such as *NFE2L2* mutations in the squamous cell subtype of esophageal cancers (Network, 2017), *TP53* mutations in Human Papillomavirus-negative tumors in head and neck cancer (Network, 2015), *KIT* mutations in testicular seminomas (Kemmer et al., 2004) and *PIK3CA* mutations in the Luminal A subtype of breast cancer (Network, 2012b). It should be noted that in some cases these differences are confounded by the fact that subtypes were originally defined by mutation status (GBM or LGG with *IDH1/IDH2* mutations).

Rare driver missense mutations exist not only in rarely mutated driver genes, but also may be spatially proximal in protein structure to common driver missense mutations. For example, the protein phosphatase PPP2R1A, which has been implicated as a tumor suppressor gene in many tumor types (Jeong et al., 2016), contained common driver missense mutations in our pan-cancer analysis at residue positions 179 and 183, which is located at the protein interface composing the phosphatase 2A complex (Figure 3d). It also had a much broader set of rare driver mutations throughout the protein interface, such as R105Q and R459C. Similarly, CHASMplus identified common driver missense mutations (S310A/F/Y) in the extracellular domain of the well-known oncogene ERBB2, but also finds rare driver missense mutations in both the extracellular and kinase domain (e.g., L313V and R678Q) (Figure 3e). This is supportive of previous experimental work implicating rare cancer driver mutations in commonly mutated cancer driver genes(Kim et al., 2016).

Truncating or likely loss-of-function mutations are typical hallmarks of tumor suppressor genes (Vogelstein et al., 2013). However, the role of driver missense mutations may be undercharacterized in tumor suppressor genes, since these mutations are more diverse and occur over a larger region than in oncogenes (Porta-Pardo et al., 2017; Tokheim et al., 2016a). As a case in point, the tumor suppressor gene *CASP8* contains many truncating variants, while all of the putative driver missense mutations identified by CHASMplus were considered rare (Figure 3f). *CASP8* is a member of the apoptosis pathway and recently has been associated with a potential role in immune evasion in cancer (Rooney et al., 2015; Thorsson et al., 2018).

We explored functional evidence to support whether the rare driver missense mutations in *CASP8*, predicted by CHASMplus, behaved similarly to truncating variants. Shmulevich and colleagues previously characterized immune-related biomarkers in TCGA tumor samples, *i.e*., inferred levels of immune cell infiltrates and expression of immune-related genes (Thorsson et al., 2018). For 12 immune-related markers, we compared tumor samples with driver missense mutations or truncating mutations in *CASP8* to control samples with no mutations in *CASP8*. In Head & Neck Squamous Cell Carcinoma (HNSC), both types of mutated samples had higher estimated levels of leukocytes, CD8 T-cells and Th1 response than controls (adjusted p≤0.001, Mann-Whitney U test, for all except truncating mutations in Th1 response, adjusted p=0.01), and significantly elevated expression of key genes involved in tumor immunity, i.e., *PDCD1* (PD-1; missense: adjusted p=1.6E-04; truncating: adjusted p=3E-02), *CD274* (PD-L1; missense: adjusted p=1E-04, truncating: adjusted p=1E-03), *CD8A* (missense: adjusted p=1E-04; truncating: adjusted p=1E-03), and *CTLA4* (missense: adjusted p=3E-03; truncating: adjusted p=4.5E-02) (Figure 4). Conventional wisdom has suggested that because rare missense mutations in tumor suppressor genes do not tend to cluster in protein sequence, they are solely passenger events (Vogelstein et al., 2013). However, our work suggests that rare driver missense mutations in *CASP8* and perhaps in other tumor suppressor genes may be relevant to tumor immune-related phenotypes. This may be partially explained by the visual appearance that distal mutations in CASP8 protein sequence show slightly greater clustering while viewed on protein structure (Figure S5). Further experimental evidence will be needed to confirm a causal relationship.

**Figure 4.**
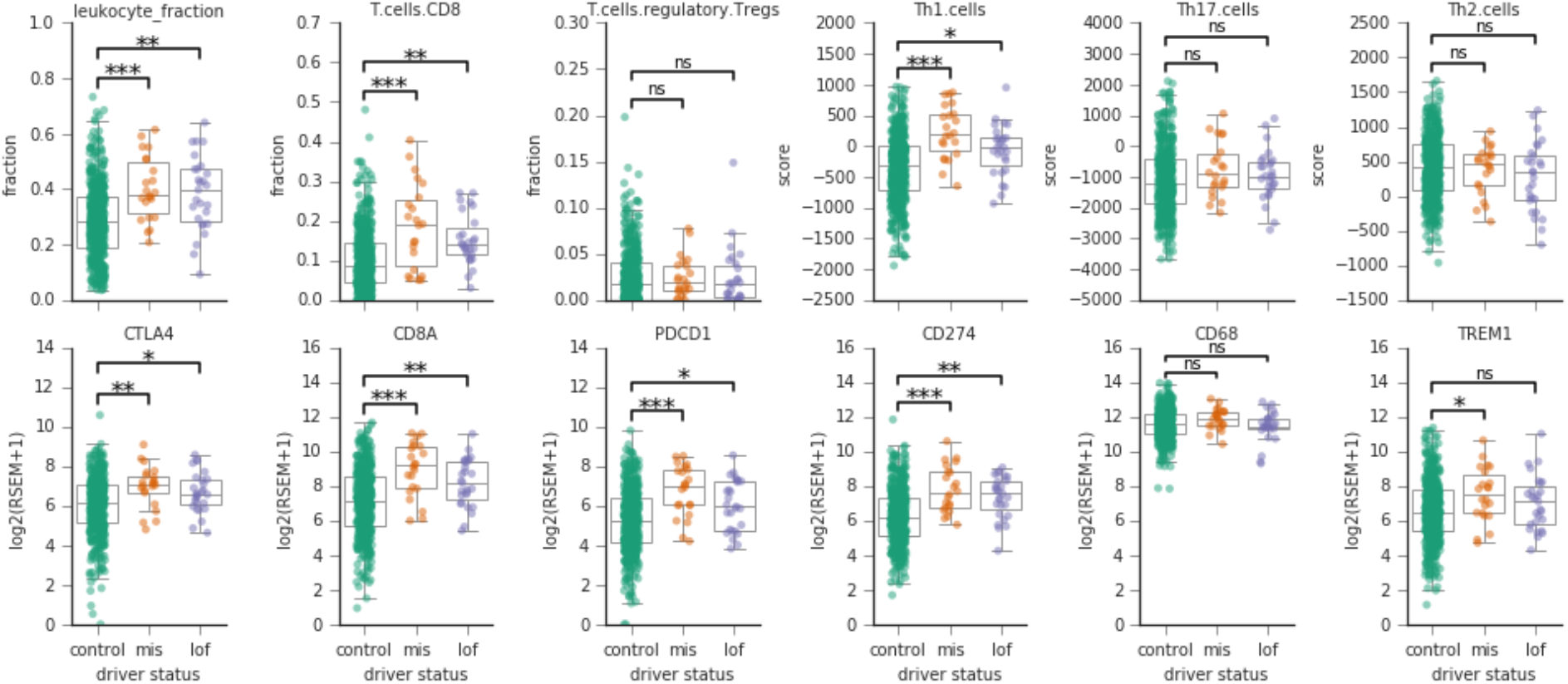
Inferred immune cell content and gene expression correlate with presence of predicted driver missense mutations in CASP8. Twelve immune-related biomarkers are shown, as estimated in Head and Neck Squamous Cell Carcinoma (HNSC) tumor samples from TCGA (Thorsson et al., 2018). Each panel compares the distribution of a marker in samples harboring a rare driver missense mutation (orange), a truncating mutation (purple) and control samples with no CASP8 mutations (green). Both tumor samples with rare driver missense mutations and truncating mutations showed a similar significantly elevated inferred immune cell infiltration. Top row, inferred immune infiltrates from DNA methylation or gene expression from TCGA HNSC samples (Thorsson et al., 2018). Bottom row, gene expression values from RNA-Seq for several important immune-related genes reported in (Thorsson et al., 2018). “mis” indicates samples with driver missense mutations identified by CHASMplus, and “lof” is likely loss-of-function variants (nonsense, frameshift insertion/deletions, splice site, translation start site, and nonstop mutations). Mann-Whitney U test, adjusted p-value (Benjamini-Hochberg method): *<0.05, **<0.01, and ***<0.001.

### CHASMplus for *in silico* saturation mutagenesis

*In silico* saturation mutagenesis can potentially complement multiplexed functional assays that assess all mutations in genes of interest, regardless of whether they have been seen before (Fowler and Fields, 2014). These experiments have so far been limited to handful of well-characterized genes and are not yet available for most genes implicated in cancer (Bailey et al., 2018). In contrast, *in silico* saturation mutagenesis can be applied on any gene, but its accuracy is unclear. We compared CHASMplus predictions (Figure 5a) to two multi-plexed functional assays characterizing the tumor suppressor gene *PTEN*. The protein product of this gene is a lipid phosphatase of phosphatidylinositol (3,4,5)-trisphosphate, an important signaling molecule in the PI3K signaling pathway, which is often dysregulated in human cancers (Sanchez-Vega et al., 2018). Because *PTEN* mutations are found in many cancer types (Bailey et al., 2018), we used CHASMplus pan-cancer predictions. The first functional assay systematically measured lipid phosphatase activity (Mighell et al., 2018), while the second measured intracellular PTEN protein abundance (Matreyek et al., 2018), potentially an indicator of thermodynamic stability. Concordant with its tumor suppressor role, driver scores from CHASMplus are negatively correlated with both lipid phosphatase activity and PTEN protein abundance (Figure 5b-c, Table S6), indicating CHASMplus was able to identify functionally damaging mutations. Interestingly, CHASMplus predictions were more correlated with each of the functional assays (lipid phosphatase activity spearman *ρ*=-0.52, protein abundance spearman *ρ*=-0.43) than they were to each other (spearman *ρ*=0.35), suggestive that it may be capturing more than one mode of damage in PTEN (Figure 5d).

**Figure 5.**
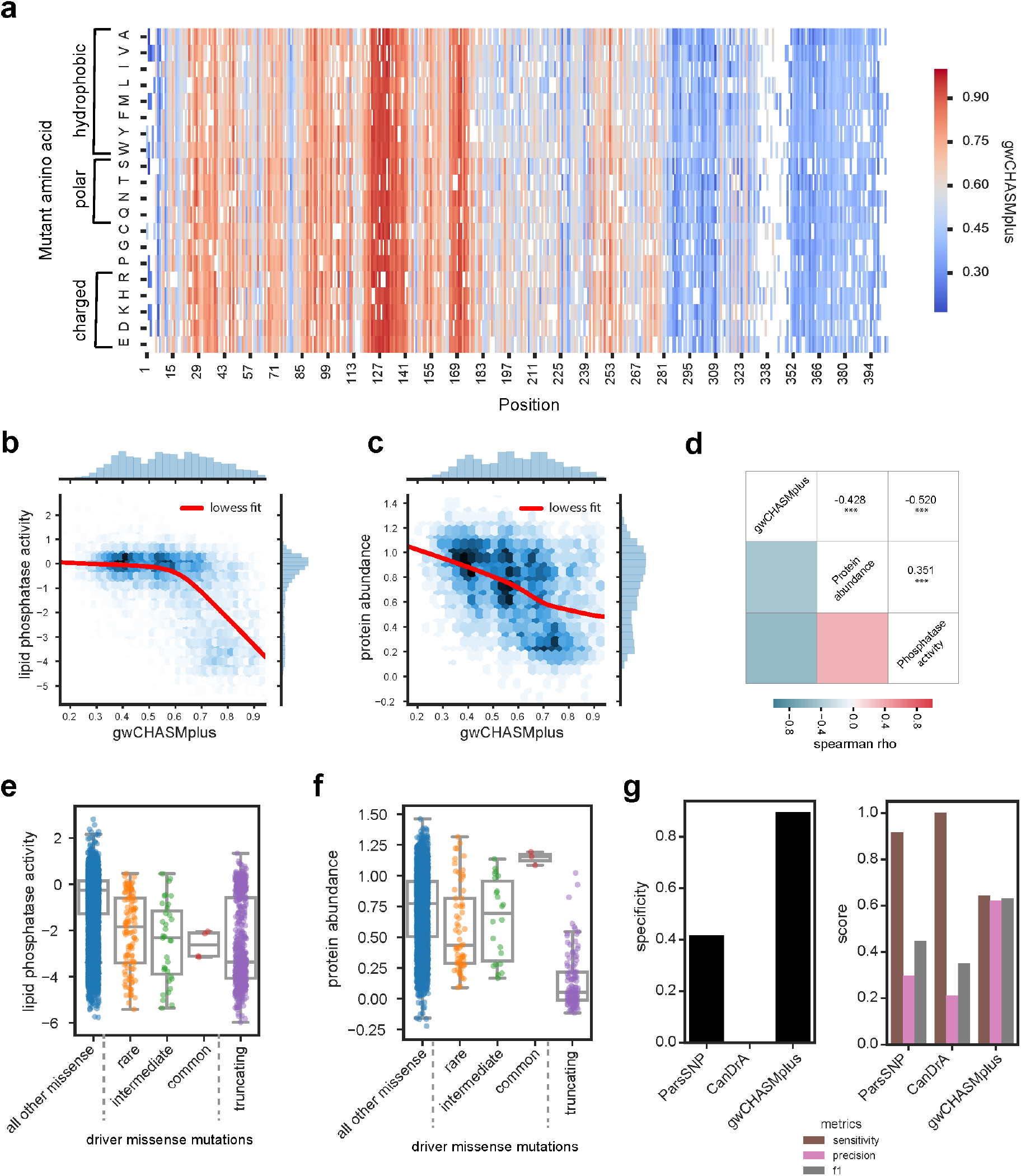
CHASMplus predictions correlate with multiplexed functional assays in PTEN. **a)** Heatmap displaying gene-weighted CHASMplus scores (gwCHASMplus) of PTEN missense mutations. **b)** Scores are negatively correlated with PTEN lipid phosphatase activity (spearman rho=-0.520) (Mighell et al., 2018) and **c)** PTEN protein abundance (spearman rho=-0.428) (Matreyek et al., 2018). **d)** Both gwCHASMplus correlations are absolutely larger than the correlation between the two experiments (spearman rho=0.351). ***=p<0.001. Distribution of **e)** PTEN lipid phosphatase activity or **f)** protein abundance in predicted driver missense mutations from TCGA (common: >5% of tumor samples, intermediate: 1-5%, and rare: <1%), all other missense mutations and truncating mutations. There is a statistically significant difference between predicted drivers and other missense mutations. The relationship between predicted driver missense and truncating mutations differs between the two assays (see Results). **g)** Comparison of CHASMplus to the 2^nd^ and 3^rd^ ranked methods in Figure 2e. Left, specificity of methods at identifying PTEN missense mutations that do not lower lipid phosphatase activity. Right, sensitivity (recall), precision, and F1 score for identifying missense mutations that lower lipid phosphatase activity. CHASMplus had the highest specificity, precision and F1 score.

Next, we examined the lipid phosphatase activity and protein abundance for the PTEN mutations that we predicted as drivers in the TCGA. We observed that these driver missense mutations, regardless of frequency, had significantly lower lipid phosphatase activity than other missense mutations in *PTEN* (common: p=0.008; intermediate: p=1.9e-9; rare: p=1.6e-18; Mann-Whitney U test, Figure 5e). Moreover, when we compared the median lipid phosphatase activity of rare, intermediate, and common driver missense mutations, we saw a trend towards lower lipid phosphatase activity. Truncating mutations had the lowest activity (p=1.6e-112, Mann-Whitney U test). A likely explanation is that greater decreases in PTEN lipid phosphatase activity may promote tumor growth and tumor clones with these mutations are positively selected in many patients. This would result in more damaging PTEN variants being more frequently observed. In contrast, we did not find a similar pattern associating increased frequency with lower PTEN protein abundance. These results are consistent with reduced lipid phosphatase activity being more correlated with variants labeled as pathogenic in ClinVar than protein abundance (Figure S6).

Next, we considered whether ParsSNP and CanDrA (2^nd^ and 3^rd^ ranked in pan-cancer benchmarks, Figure 2e) could predict the lipid phosphatase activity of PTEN. Compared to these methods, CHASMplus demonstrated substantially higher specificity (specificity=0.89 vs. 0.42 and 0.0, ParsSNP and CanDrA, respectively), and the difference is statistically significant (p<2.2e-16, McNemar’s test, Figure 5g). Specificity is defined as the proportion of negative examples, in this case lack of damage to lipid phosphatase activity, that are correctly predicted. Next, we applied the metrics of sensitivity, precision and F1 score (harmonic mean of sensitivity and precision) (Figure 5g). CHASMplus had the best balance of sensitivity and precision (highest f1 score) and the best precision. The other two methods have higher sensitivity at identifying damaging mutations, but at the cost of poor specificity. We attribute the better performance of CHASMplus to our control of errors using the statistically rigorous false discovery rate, which is not used by the other methods.

### The trajectory of driver discovery across diverse cancer types

We next sought to understand whether cancer types fundamentally differed in their usage of driver missense mutations. Based on our cancer type-specific models from CHASMplus, we found that the diversity and prevalence of driver missense mutations varied considerably across TCGA cancer types (Figure 6a, Methods). Diversity was defined with respect to the distribution of driver missense mutations across codons and prevalence with respect to the overall frequency of mutations in tumor samples. High diversity mutations are broadly distributed across codons, while high prevalence mutations occur in a large proportion of tumor samples. Using K-means clustering, we found that cancer types could be grouped into high diversity and low prevalence (12 cancer types), high diversity and high prevalence (15 cancer types), and low diversity and high prevalence (5 cancer types). These differences were not associated with intra-tumor heterogeneity or normal contamination, as assessed by mean variant allele fraction (VAF) of a cancer type (p>0.05, correlation test, Methods). Nor were they associated with TCGA sample size for a particular cancer type. For example, while both pancreatic ductal adenocarcinoma (PAAD) and sarcoma (SARC) had similar sample sizes (n=155, n=204 respectively), PAAD had high prevalence and low diversity, while SARC had low prevalence and high diversity.

**Figure 6.**
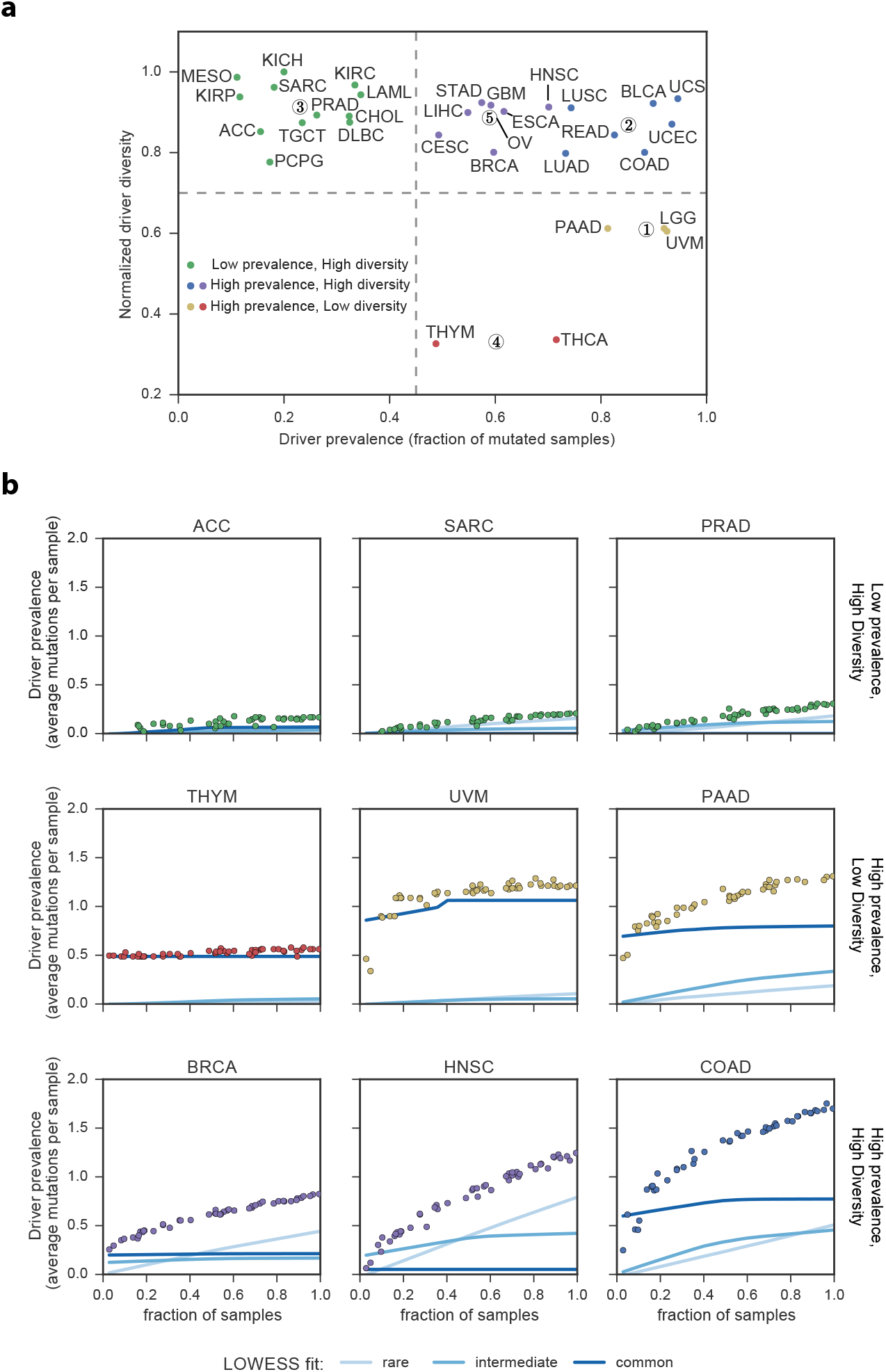
Characteristics and trajectory of missense mutation driver discovery. **a)** Plot displaying normalized driver diversity and driver prevalence (fraction of tumor samples mutated) for driver missense mutations in 32 cancer types. K-means clustering identified 5 clusters with centroids shown as numerically designated circles. **b)** Prevalence of driver missense mutations identified by CHASMplus as a function of sample size. Lines represent LOWESS fit to different rarities of driver missense mutations. TCGA acronyms for cancer types are listed in the Methods.

Are there substantially more cancer driver missense mutations yet to be discovered? Prior attempts to address this question have focused on the identification of cancer driver genes (Davoli et al., 2013; Lawrence et al., 2014; Martincorena et al., 2017), which can contain a confounding mixture of both passenger and driver mutations. Here we examined the trajectory of driver discovery from CHASMplus at the *mutation-level* as the number of tumor samples analyzed is gradually increased by random subsampling. Subsampling analysis showed all cancer types had a linear increase in the number of unique driver missense mutations identified (*R*^2^ > 0.5) with no evidence of saturation at current sample sizes (Figure S7a). However, discovery of driver missense mutations, which occur in aggregate at a given prevalence (average number per cancer sample), varied substantially among cancer types (Figure 6b). For sarcoma (SARC), adrenocortical carcinoma (ACC), and prostate adenocarcinoma (PRAD), as sample size increased there was minimal increase in the prevalence of driver missense mutations. As a case in point, we extended our analysis to data from a recently released PRAD study (Armenia et al., 2018), which augmented the 477 TCGA PRAD samples with 536 additional samples. This resulted in only marginal increases in the overall prevalence of identified driver missense mutations, consistent with our predicted trajectory based only on TCGA samples (Methods, Table S7, Figure S7b). In contrast, thymoma (THYM), uveal melanoma (UVM), and pancreatic ductal adenocarcinoma (PAAD) contained common driver missense mutations that could be detected based on only a few samples from the cohort, *e.g*., GTF2I L424H in THYM. Prevalence of driver missense mutations abruptly saturated for THYM and UVM as sample size increased, and nearly all of these mutations were common. In PAAD the overall driver prevalence exhibited a diminishing rate of discovery, but the prevalence of intermediate or rare driver missense mutations increased with greater sample size. In contrast, the prevalence of rare driver missense mutations increased substantially with sample size in breast cancer (BRCA), head and neck squamous cell carcinoma (HNSC), and colon adenocarcinoma (COAD).

These results suggest cancer types can be clustered by patterns of driver missense mutation diversity and prevalence (Figure 6a), in addition to well-established approaches to define cancer subtypes, such as by cell-of-origin (Hoadley et al., 2018). Moreover, a statistical power analysis suggests that an alternative approach based only on mutation hotspot detection would be underpowered to detect such results (Figure 7, Methods).

**Figure 7.**
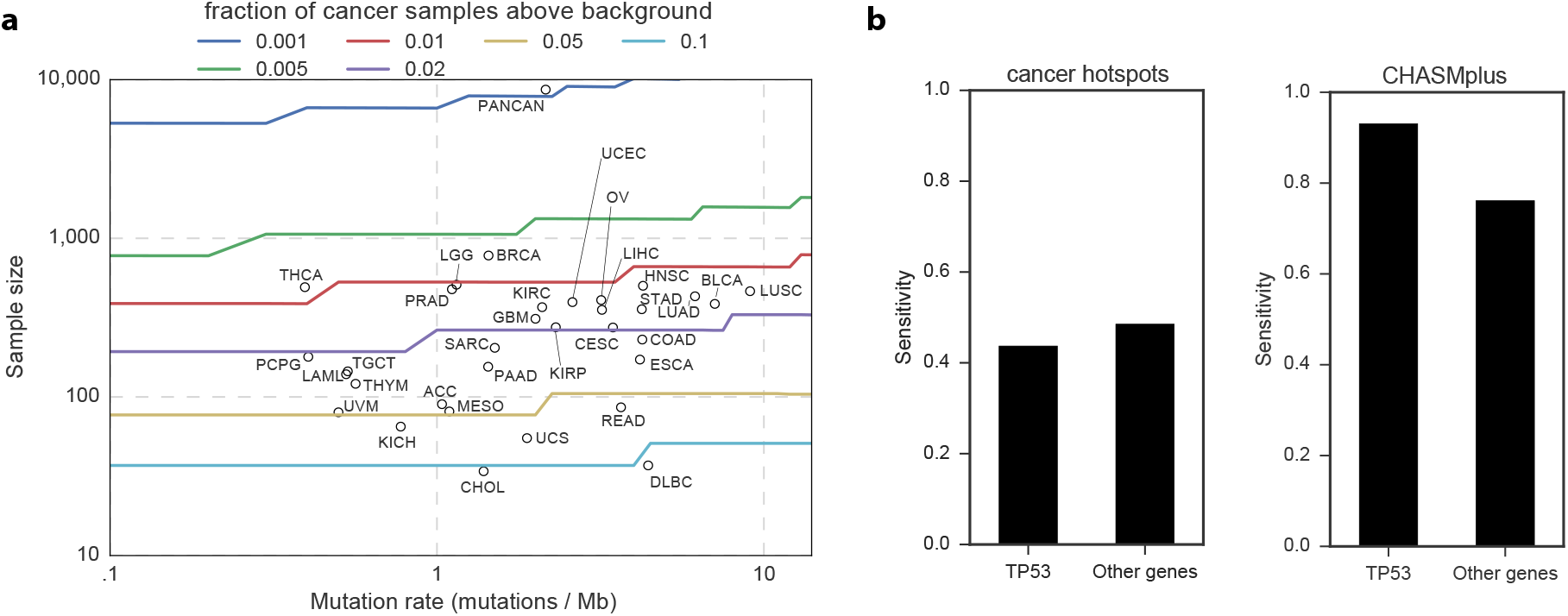
Hotspot detection alone has limited statistical power to identify driver mutations. **a)** Statistical power to detect a significantly elevated number of non-silent mutations in an individual codon, as a function of sample size and mutation rate. Circles represent each cancer type from the TCGA and are placed by sample size and median mutation rate. Curves are colored by the frequency of driver mutations (fraction of non-silent mutated cancer samples above the expected background mutation rate). If a circle is below a curve, then hotspot detection is not yet sufficiently powerful to detect driver mutations of that frequency. **b)** Bar graph comparing power (sensitivity) to detect labeled oncogenic driver missense mutations from OncoKB between CHASMplus and the cancer hotspots method (Chang et al., 2016). Stratification by *TP53* suggests that the increased power provided by CHASMplus is not solely a result of high performance on oncogenic *TP53* mutations.

## Discussion

Cancer genome interpretation is challenging due to the reality that of all somatic mutations observed in cancer, only a small proportion are drivers (Tomasetti et al., 2015). Future insights into cancer evolution and its relevance for clinical care will increasingly rely on the precise interpretation of whether individual mutations are cancer drivers(Hyman et al., 2017b). Currently available computational methods to support driver mutation discovery need to be improved to increase their clinical relevance (Martelotto et al., 2014; Molina-Vila et al., 2014). CHASMplus is the first computational method that can predict driver missense mutations for specific cancer types, an important advance in the field, as a patient’s therapeutic response to drugs targeting a specific gene and optimal assignment to a clinical trial is increasingly understood to depend on both the specific mutation in the gene of interest and cancer type (Brose et al., 2016; Hyman et al., 2018; Hyman et al., 2017a; Prahallad et al., 2012). We are the first to use semi-supervised learning to this problem (Methods). On both cancer type-specific and pan-cancer benchmarks, CHASMplus had significantly higher performance than comparable methods.

The long tail hypothesis(Ding et al., 2010; Garraway and Lander, 2013) posits that there are many rare driver mutations in human cancers. To explore this hypothesis, we leveraged the improvements made in CHASMplus to systematically predict driver missense mutations in 8,657 cancer samples from the TCGA. We predict that rare driver missense mutations as a group are common in cancer. However, not every type of cancer is the same; our study is the first, to our knowledge, to systematically show that the prevalence of rare driver missense mutations is highly variable across cancer types. The observed diversity of driver mutations across patients’ tumors may be influenced by tumor mutation burden, the type of gene (*i.e*., tumor suppressor genes), the functional strength of the mutation, and the mutation’s subtype specificity. Other factors may also contribute, such as epistasis (Kent et al., 2015; Skoulidis et al., 2015), interactions with the (micro)environment (Marty et al., 2017; Rooney et al., 2015), selection by cell-of-origin (Bailey et al., 2018), competition from other types of somatic alterations, or population-level differences (Paez et al., 2004). The diversity of driver missense mutations supports the critical role of understanding the origins and overall contribution of rare driver mutations -- failure to capture and identify rare driver mutations, which occur in aggregate at reasonable prevalence, may result in crucial missed opportunities for interpreting a patient’s cancer.

Because, individually, they are infrequent, rare driver mutations in newly sequenced tumors may not have been previously observed in the TCGA or other large-scale sequencing projects. One approach to characterize such singleton mutations is to run multiplexed functional assays, which functionally assess all mutations in genes of interest, regardless of whether they have been seen before (Fowler and Fields, 2014). These experiments have so far been limited to handful of well-characterized genes and are not yet available for most genes implicated in cancer (Bailey et al., 2018). Consequently, improved computational methods are needed to prioritize mutations for low- and medium- throughput studies. We therefore have precomputed the score of every possible missense mutation across the genome, effectively an *in silico* saturation mutagenesis across all genes to score as of yet unobserved mutations that are potential cancer drivers.

There are several limitations to our study. Although missense mutations are the most frequent protein-coding somatic alteration in cancer (Vogelstein et al., 2013), CHASMplus only predicts missense mutations; however, in principle, our approach could be extended to other types of alterations. Additionally, although the scope of our study focused on driver mutations, passenger mutations may also have clinical relevance, such as by forming neoantigens that mediate response to immune checkpoint blockade therapy (Schumacher and Schreiber, 2015). Lastly, while our study leverages the large sized TCGA cohorts to make informed judgements about driver missense mutations, inevitably these tumors may still miss important contexts for understanding cancer. The collection of additional tumor samples especially from understudied populations or cancer subtypes could yield unexpected improvements that are not currently captured. For example, it is well known *EGFR* mutations in lung adenocarcinoma are more highly prevalent in Asian compared to Caucasian populations (Paez et al., 2004). However, this is a general limitation for all cancer sequencing studies (Bailey et al., 2018; Davoli et al., 2013; Lawrence et al., 2014; Lawrence et al., 2013; Martincorena et al., 2017) and is not specific to CHASMplus.

We expect that an increasingly complete catalog of driver missense mutations will be generated by a combination of improved computational methods and cumulative growth of available samples from cancer sequencing studies. However, for some cancer types in some populations, discovery of driver missense mutations may already be effectively saturated. Studies based on driver genes have predicted a linear increase in discovery with increasing sample size (Armenia et al., 2018; Davoli et al., 2013; Lawrence et al., 2014). By analyzing the trajectory of discovery at the level of driver missense mutations, we identified a more complicated pattern which depends on the cancer type. Increases predicted by gene-level trajectories may be inflated because both driver and passenger mutations are contributing to prevalence. Future work will further elucidate a broader range of driver mutations, including those within non-coding regions of the genome, at different stages of carcinogenesis, such as in pre-cancerous lesions, and in response to therapeutic treatment.

### Availability

TCGA mutation scores (http://chasmplus.readthedocs.io/). Interactive portal (http://karchinlab.github.io/CHASMplus).

## Acknowledgements

*Funding:* National Cancer Institute (NCI) Grant F31CA200266 (to C.T.); NCI Grant U24CA204817 (to R.K.)

## Author Contributions

CT and RK conceived of the study. CT and RK drafted and edited the manuscript. C.T. developed the CHASMplus algorithm and analyzed results.

## Supplemental Figure Legends

**Supplementary Figure 1.**
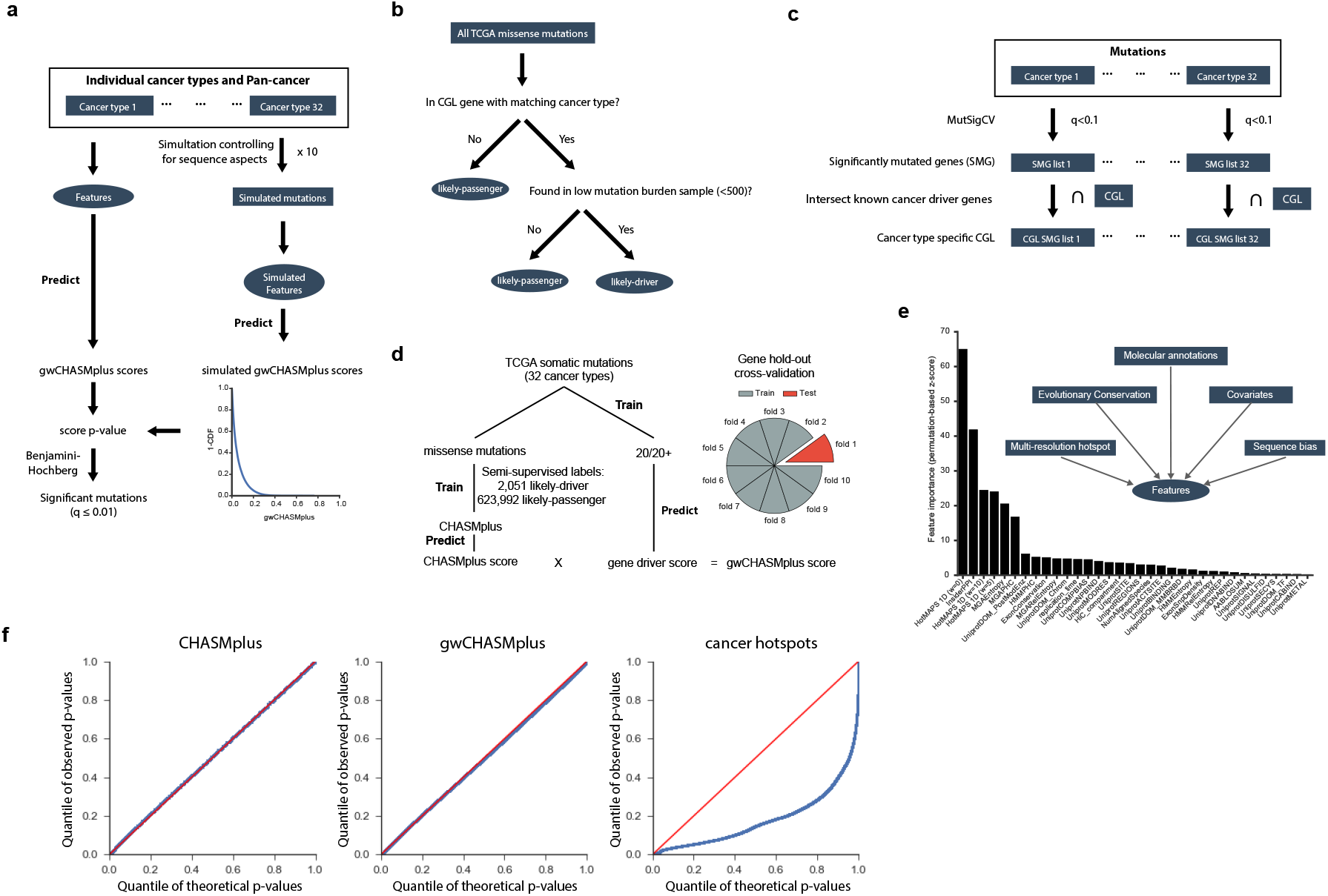
Overview of CHASMplus. Related to Figure 1. **a)** Diagram of how CHASMplus identifies driver missense mutations based on models identifying mutations with statistically significant scores. CHASMplus models are trained using a semi-supervised approach. **b)** Missense mutations were labeled either as “likely-passenger” or “likely-driver” based on a two-step approach: overlap with previously known genes from Cancer Genome Landscapes (CGL) in a cancer type-specific manner and tumor samples with low mutation burden. **c)** Diagram demonstrating how the cancer type-specificity of CGL genes were determined based on MutSigCV. **d)** Training and testing procedure for CHASMplus. The gene hold-out cross-validation was applied to both subsequent driver discovery and performance benchmarking efforts. **e)** Features with a net-positive feature importance according to a permutation adjusted z-score are shown. Boxed text indicates broad feature categories that were important. **f)** CHASMplus p-values are well calibrated as evidenced by a QQ plot of observed p-values (blue line) compared to theoretically expected under the null hypothesis (red line). CHASMplus represents unweighted CHASMplus scores, gwCHASMplus represents gene weighted CHASMplus scores, and cancer hotspots is a codon-level hotspot detection method (Chang et al., 2016). All mutations in genes found in the Cancer Gene Census were removed to eliminate possible driver mutations in this comparison, and consequently the remaining mutations will have greater enrichment for passenger mutations (null hypothesis) in the QQ plot.

**Supplementary Figure 2.**
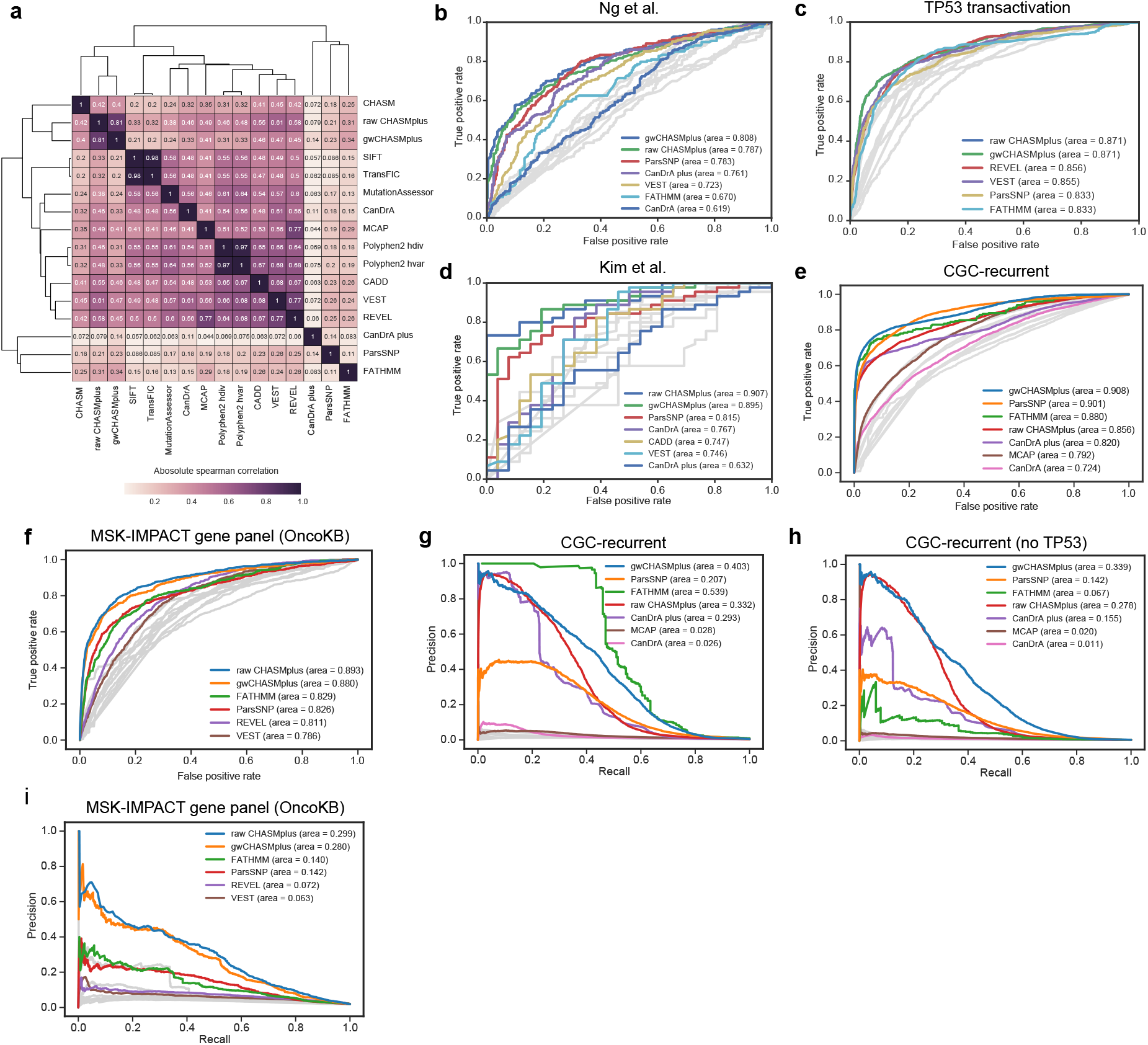
Detailed pan-cancer benchmark performance. Related to Figure 2. **a)** Heatmap of the absolute spearman correlation between methods on the TCGA mutation dataset. **b-f)** Receiver Operating Characteristic curves for, in order, the following benchmarks: Ng *et al., TP53* transactivation (IARC database), Kim et al., recurrent mutations in the Cancer Gene Census, and annotated oncogenic mutations using OncoKB on the MSK-IMPACT gene panel. Area under the curve is shown in parenthesis. Top 5 methods are labeled, but if the method has two version of scores then both are shown. Precision-recall curve for imbalanced benchmarks: **g)** recurrent mutations in the Cancer Gene Census (CGC-recurrent); **h)** To identify potential overfitting by methods, we repeated the Precision-Recall curve with all TP53 mutations removed; **i)** MSK-IMPACT gene panel using OncoKB. Differential performance between panels g and h is a reasonable indicator of overfitting for a method, suggesting FATHMM and CanDrA plus may have overfit to *TP53*. Area under the curve is shown in parenthesis. Top 5 methods are labeled according to auROC performance, but if the method has two version of scores then both are shown.

**Supplementary Figure 3.**
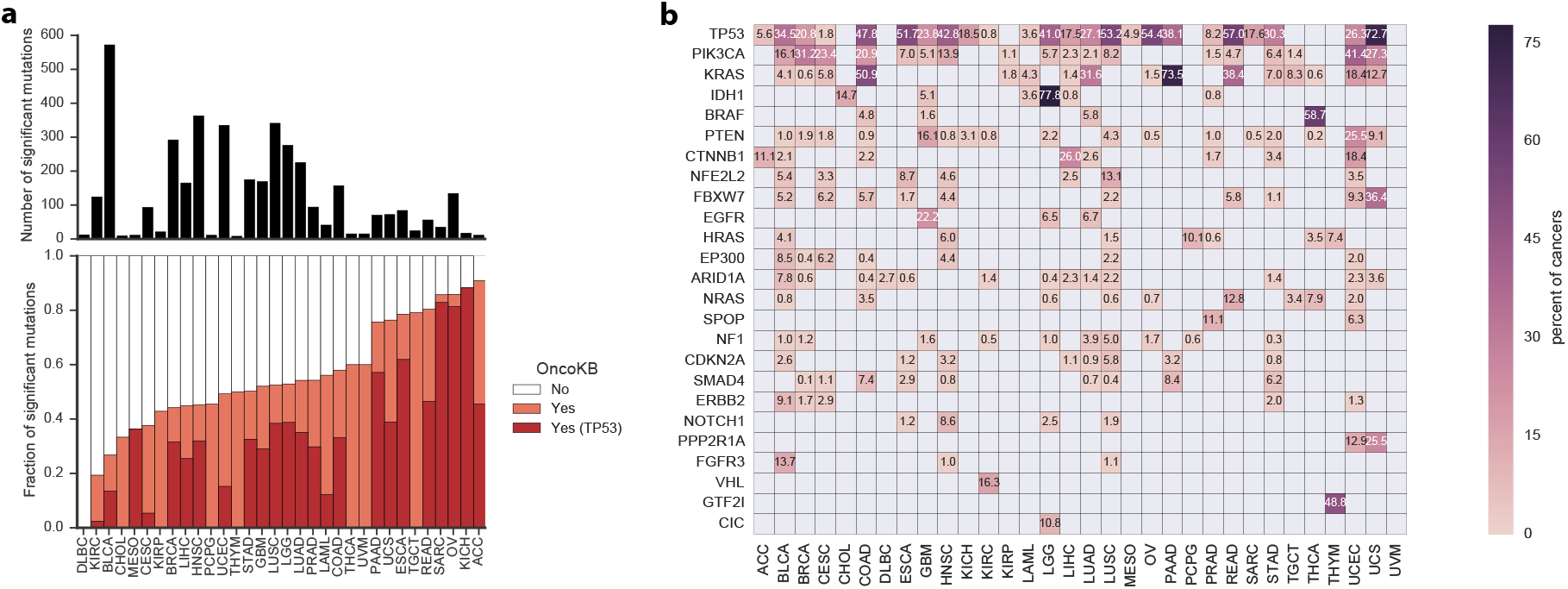
Driver missense mutations found by CHASMplus that do not appear in OncoKB. Related to Figure 3. CHASMplus identified putative driver missense mutations that are not in OncoKB, a highly-curated database of oncogenic mutations. **a)** Bar graphs showing of the number of unique driver missense mutations by cancer type (top) and the proportion that appear in OncoKB (bottom). **b)** Top 25 genes for each cancer type with the most frequent driver missense mutations in TCGA. Heatmap colored by percentage of samples that are mutated.

**Supplementary Figure 4.**
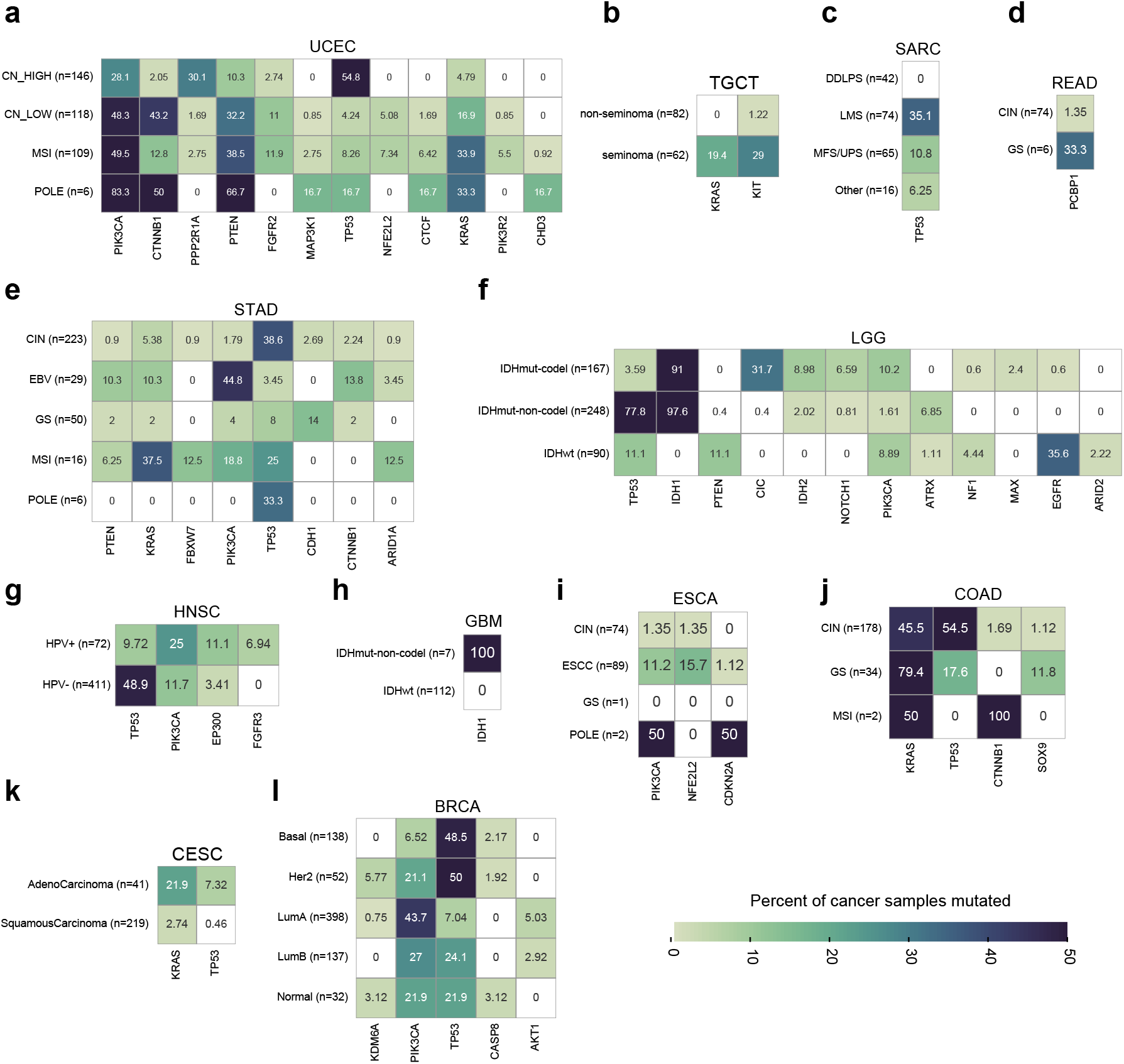
Subtype enrichment for driver missense mutations at the gene-level. Related to Figure 3. All genes with a statistically significant preferential enrichment for driver missense mutations in one or more cancer subtypes are shown in the form of a heatmap (q<0.1, chi-squared test). Heatmaps are formatted as follows: the cancer type is noted above the heatmap, the y-axis represents cancer subtypes, and the x-axis represents genes. The percentage of samples containing driver missense mutations is indicated in each heatmap cell. **a-l)** Heatmap results, in order, for UCEC, TGCT, SARC, READ, STAD, LGG, HNSC, GBM, ESCA, COAD, CESC, and BRCA. Cancer subtype information was obtained from (Sanchez-Vega et al., 2018).

**Supplementary Figure 5.**
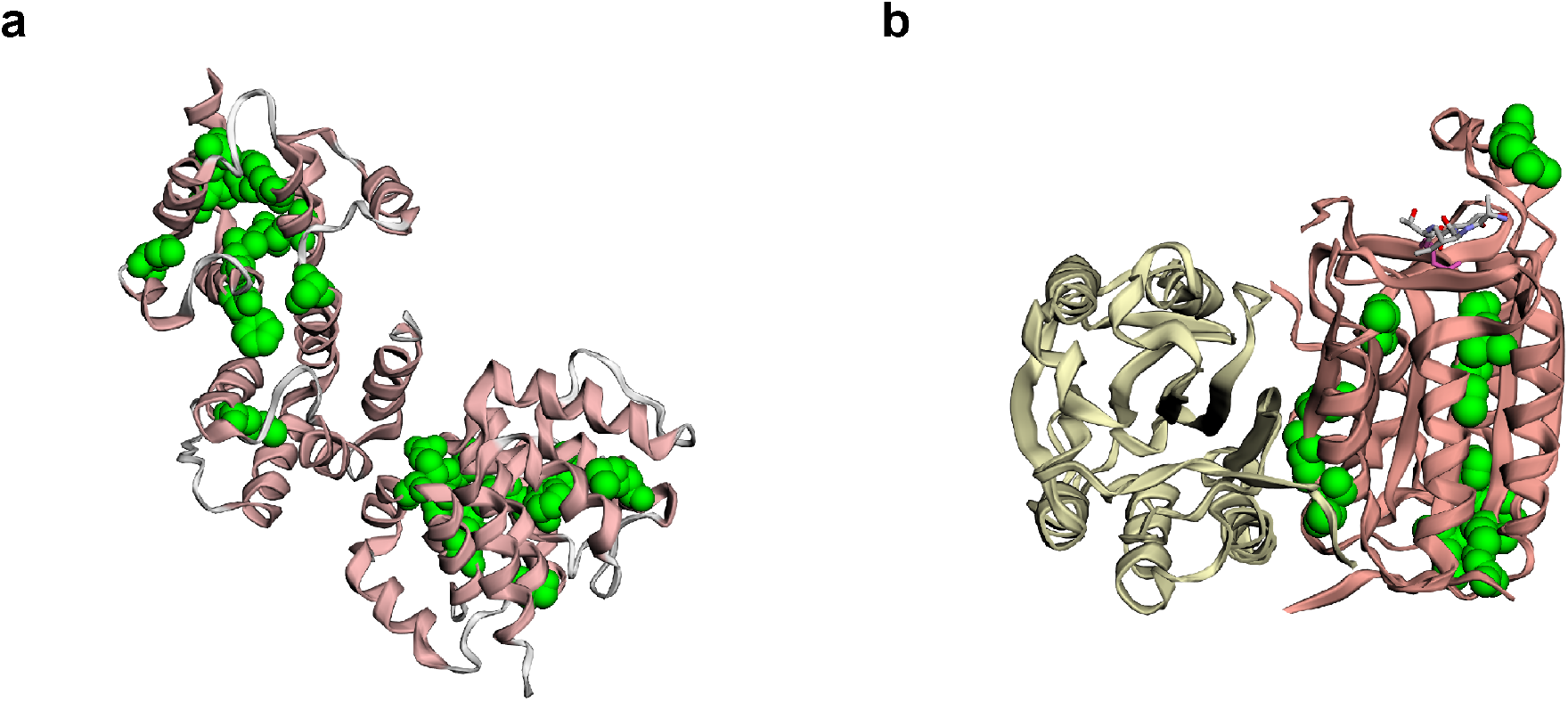
Localization of predicted driver missense mutations in CASP8 protein structure. Related to Figure 4. **a)** Protein structure of tandem Death Effector Domains of CASP8 (PDB: 4ZBW). **b)** Protein structure of the peptidase-like domain of CASP8 (salmon) and a CASP8-regulator c-FLIPL (white) (PDB: 4ZBW).

**Supplementary Figure 6.**
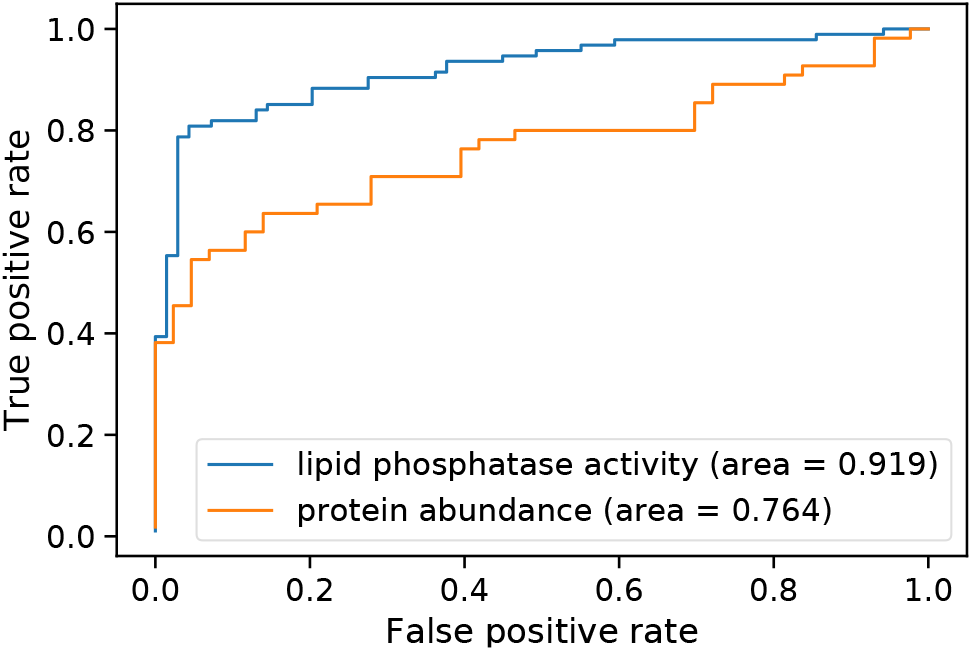
Performance of multiplexed functional assays at discriminating ClinVar pathogenic mutations in PTEN. Related to Figure 5. The functional scores derived from two studies, which measure PTEN lipid phosphatase activity (blue, (Mighell et al., 2018)) and protein abundance (orange, (Matreyek et al., 2018)), were compared to pathogenicity annotations from ClinVar. Performance was measured by the area under the Receiver Operating Characteristic curve.

**Supplementary Figure 7.**
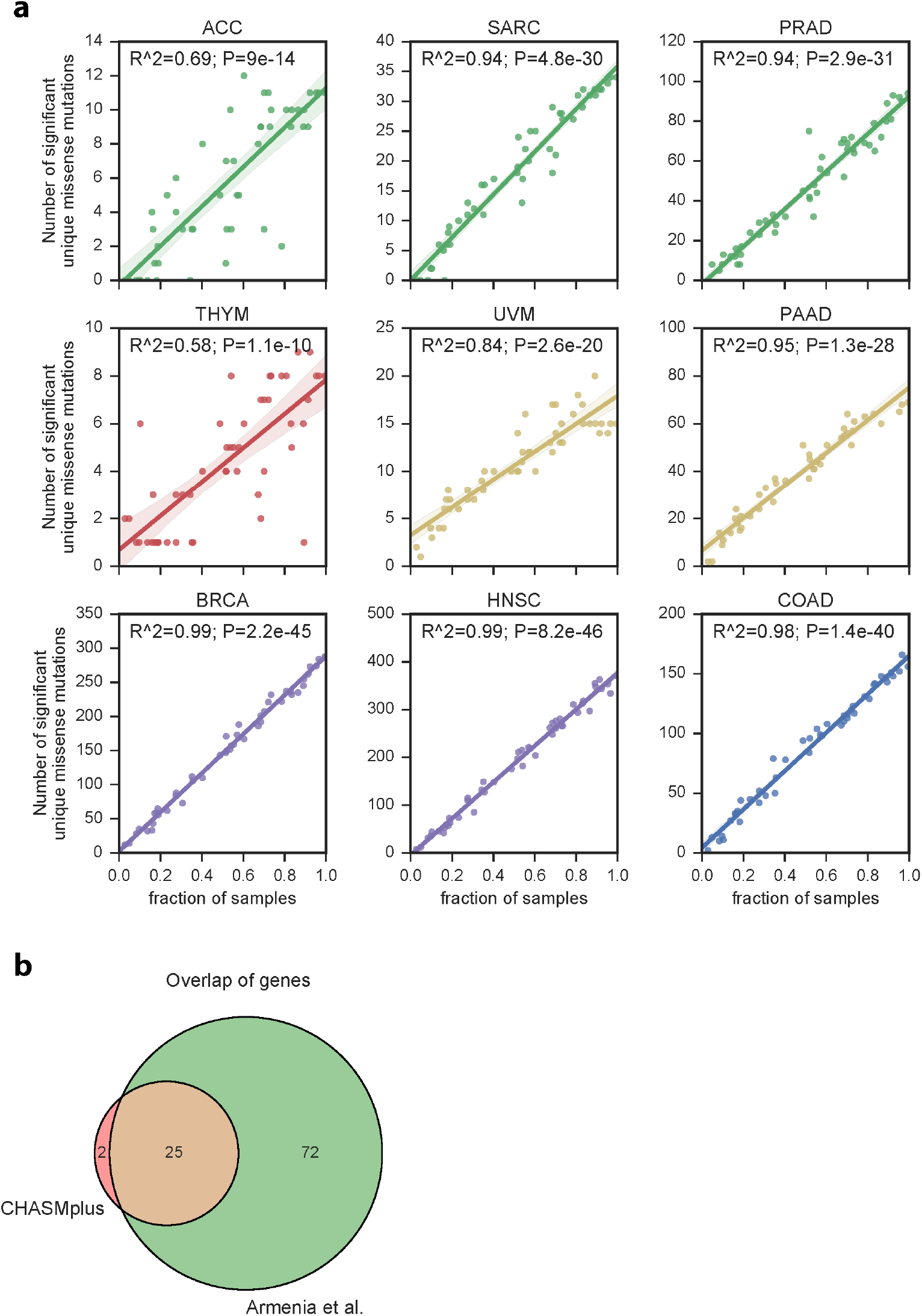
Trajectory analysis of putative driver missense mutations in nine cancer types as sample size increases. Related to Figure 6. CHASMplus was run on resampled subsets of the TCGA mutation dataset for each cancer type, as the number of tumor samples was increased. **a)** Number identified as significant by CHASMplus (q<=0.01) as a function of sample size. Unique driver missense mutations show a linear increase, but overall prevalence has a sub-linear increase (Figure 6b). **b)** Evidence from a recently published prostate adenocarcinoma (PRAD) sequencing study that was larger than the TCGA supports our predicted sub-linear increase in driver prevalence with increased sample size (Methods).

## STAR Methods

### CONTACT FOR REAGENT AND RESOURCE SHARING

For additional information regarding the data, please contact Rachel Karchin.

### EXPERIMENTAL MODEL AND SUBJECT DETAILS

The Cancer Genome Atlas (TCGA) used informed consent to collect human samples with approval of local institutional review boards (https://cancergenome.nih.gov/abouttcga/policies/informedconsent).

### TCGA Mutation dataset

We collected a set of 1,225,917 somatic mutations in 8,657 samples from The Cancer Genome Atlas (TCGA) somatic mutation calls from whole-exome sequencing (v0.2.8, https://synapse.org/MC3)(Ellrott et al., 2018). We analyzed 32 cancer types, with abbreviations for the cancer types listed below. We further filtered mutations by restricting to only mutations annotated with a ‘PASS’ filter, except for OV and LAML where mutations with only whole genome amplified (‘wga’) status were allowed because otherwise the majority of samples would be filtered. We additionally removed hypermutated samples, as they tend to have an adverse effect on statistical power. We identified hypermutated samples as having more mutations than 1.5 times the interquartile range above the third quartile (Tukey’s condition) of samples within the same cancer type. Because some relatively low mutation rate cancer types contained outliers, we additionally required the sample to have at least 1,000 mutations to be considered hypermutated.

Cancer types in the TCGA are abbreviated as follows: Acute Myeloid Leukemia (LAML, n=139), Adrenocortical carcinoma (ACC, n=90), Bladder Urothelial Carcinoma (BLCA, n=386), Brain Lower Grade Glioma (LGG, n=510), Breast invasive carcinoma (BRCA, n=779), Cervical squamous cell carcinoma and endocervical adenocarcinoma (CESC, n=274), Cholangiocarcinoma (CHOL, n=34), Colon adenocarcinoma (COAD, n=230), Esophageal carcinoma (ESCA, n=172), Glioblastoma multiforme (GBM, n=311), Head and Neck squamous cell carcinoma (HNSC, n=502), Kidney Chromophobe (KICH, n=65), Kidney renal clear cell carcinoma (KIRC, n=368), Kidney renal papillary cell carcinoma (KIRP, n=275), Liver hepatocellular carcinoma (LIHC, n=354), Lung adenocarcinoma (LUAD, n=431), Lung squamous cell carcinoma (LUSC, n=464), Lymphoid Neoplasm Diffuse Large B-cell Lymphoma (DLBC, n=37), Mesothelioma (MESO, n=81), Ovarian serous cystadenocarcinoma (OV, n=408), Pancreatic adenocarcinoma (PAAD, n=155), Pheochromocytoma and Paraganglioma (PCPG, n=179), Prostate adenocarcinoma (PRAD, n=477), Rectum adenocarcinoma (READ, n=86), Sarcoma (SARC, n=204), Stomach adenocarcinoma (STAD, n=357), Testicular Germ Cell Tumors (TGCT, n=145), Thymoma (THYM, n=121), Thyroid carcinoma (THCA, n=492), Uterine Carcinosarcoma (UCS, n=55), Uterine Corpus Endometrial Carcinoma (UCEC, n=396), Uveal Melanoma (UVM, n=80), and Pan-cancer (all cancer types) (PANCAN, n=8657).

### CHASMplus

Previous machine learning approaches for predicting driver mutations have been trained on a small positive class of bona fide driver missense mutations, which are highly prevalent in many cancer types. In contrast, we trained a model for each of 32 cancer types from TCGA with a semisupervised approach (Figure S1). Driver genes have been labeled but the labels of missense mutations are inferred by cluster assumptions, *e.g*., driver missense mutations may occur together in known driver genes and in significantly mutated genes for a particular cancer type. Mutations are assigned to the ‘positive’ class (driver-like) for a cancer type based on these assumptions, and all other mutations are assigned to the ‘negative’ class (passenger-like). To extend this approach to a pan-cancer prediction, all mutations from the positive class were merged into one group and a single classifier was trained. The availability of sequencing data from ~9,000 cancer patients covering 32 cancer types from the TCGA makes this feasible for the first time. To test the assumptions of our training procedure, we rigorously evaluated CHASMplus on orthogonally labeled mutations based on *in vitro* experiments, *in vivo* experiments, and literature curation. On both cancer type-specific and pan-cancer benchmarks, CHASMplus had significantly higher performance than comparable methods.

The code for CHASMplus is available on github (https://github.com/KarchinLab/CHASMplus).

#### Training Set

Using the TCGA mutation dataset, we established training labels with a semi-supervised approach (Figure S1b-c), designed to minimize bias. The positive class (likely-driver missense mutations) was selected by the following criteria: 1) missense mutations had to occur in a curated set of 125 pan-cancer driver genes(Vogelstein et al., 2013); 2) for each of the 32 TCGA cancer types, missense mutations found in that cancer type had to occur in a significantly mutated gene for that cancer type according to MutSigCV v1.4(Lawrence et al., 2014). We ran MutSigCV using recommended settings and a full sequencing coverage file (http://archive.broadinstitute.org/cancer/cga/mutsig). Importantly, MutSigCV v1.4 only assess the total number of mutations in a gene, and not any characteristics of those mutations; thus, we avoid making strong assumptions about the properties of a particular driver mutation; 3) missense mutations had to occur in samples with relatively low mutation rate (less than 500 mutations, half the minimum hypermutator threshold as defined above). This filter was intended to limit the number of passenger mutations mislabeled as drivers. The negative class (likely-passenger missense mutations) consisted of the remaining missense mutations in the TCGA mutation set.

For training purposes, we only used unique mutations to avoid double counting a mutation seen more than once. If, however, the same mutation had contradictory labels in different cancer types, we labeled it as a driver because it was recurrent. This established a set of 2,051 likely-driver missense mutations and 623,992 likely-passenger missense mutations, for which we found sufficient annotation to compute our selected features (Figure S1d). Skin cutaneous melanoma mutations were not included in training due to the systematically high mutation burden for this cancer type. Instead, predictions for melanoma were based on a model trained on the other cancer types (Table S3).

#### Features

CHASMplus uses a total of 95 features (Table S1), of which 34 had a net-positive feature importance as shown in Figure S1e. Important features assessed five broad categories: multiresolution missense mutation hotspots (HotMAPS 1D algorithm(Tokheim et al., 2016a)), evolutionary conservation/human germline variation, molecular function annotations (e.g., protein-protein interface annotations from(Meyer et al., 2018)), sequence biased regions, and gene-level covariates (e.g., replication timing). Interestingly, the most highly-ranked features were clustering of missense mutations in ‘hotspots’ and localization of mutations at protein-protein interfaces. Because important features such as hotspot detection and gene weights are calculated based on tumor sequencing data of a relevant cancer type, CHASMplus driver predictions can be honed for each cancer type. A broad tail in the feature importance distribution (Figure S1e) suggests that a synergy of many features also contributes to prediction performance.

#### Feature importance

We used the Mean Decrease in Gini Index (MDGI) as a measure of feature importance in the random forest. This measurement, however, has been previously noted to favor continuous features over discrete features with a small number of possible values (Altmann et al., 2010). We compensated by calculating an adjusted z-score using a permutation-based approach. This involved calculating MDGI for each feature for 1,000 permutations of mutation labels and calculating the z-score of the observed data by using the mean and standard deviation of the permutations. Permutations were designed to preserve the observed clustering of labeled driver mutations in a small number of genes, by stratifying the permutation procedure by gene. First, we randomly permuted the list of unique genes containing any missense mutation. Second, we assigned the first gene in the permuted list to have the same fraction of driver/passenger mutations as an actual gene with labeled driver mutations in our training set. We then proceeded to the next gene in the list, and repeated the procedure, until the same number of labeled driver mutations as in our actual training set was reached. All other genes had their mutations labeled as passengers. Finally, we computed the MDGI for each feature based on the CHASMplus model on the permuted training data. Grouping by gene in the permutation was done to mimic the heuristic on how training data was labeled, and to avoid gene-level features from having artificially high feature importance.

#### 20/20+ driver gene prediction

The driver gene predictions by 20/20+ (v1.2.0, https://github.com/KarchinLab/2020plus) were carried out with a Random Forest model as previously described (Tokheim et al., 2016b). We computed driver gene scores for each cancer type and pan-cancer. The driver gene score represents the fraction of decision trees predicting driver for a particular gene in the random forest.

#### Machine learning prediction

We used random forests (Amit and Geman, 1997; Breiman, 2001), a machine learning technique, to predict whether a missense mutation is a cancer driver. We trained a random forest using the *randomForest* R package.

To handle the problematic imbalance in the training set (see Training Set section), we used a stratified down sampling approach within the bagging procedure of the Random Forest. Random undersampling has been previously recommended for Random Forests based on empirical performance (Hulse et al., 2007). The imbalance occurred on two levels, there were substantially more labeled passenger missense mutations than drivers, and driver mutations were concentrated in a few well-known genes, such as *TP53*. We first calculated the median number of labeled driver missense mutations within genes containing at least one driver missense mutation label. If a gene contained more labeled driver missense mutations than the median, we set the number of driver missense mutations sampled from that gene to the median. Passenger missense mutations were sampled at an equal frequency as driver missense mutations after the gene-based median correction.

Since missense mutations in the same gene may have overlapping feature representations which result in classifier overfitting (Capriotti and Altman, 2011), we performed prediction using a 10-fold gene hold-out cross-validation procedure for both CHASMplus and 20/20+. This involved creating 10 random folds for cross-validation but ensuring all mutations within a gene were within a single fold (Figure S1d). This cross-validation protocol was applied to both our subsequent performance benchmarks and applications to TCGA and an additional prostate cancer study (Armenia et al., 2018). This ensured that predictions for a mutation were never based on a model trained on that mutation.

The CHASMplus score represents the fraction of decision trees which vote for the mutation being a driver. We calculate the gene-weighted CHASMplus score (gwCHASMplus) by multiplying the random forest score of CHASMplus by the driver gene score from 20/20+.

#### Estimation of statistical significance

Missense mutations are only considered putative drivers if scores reach statistical significance, with respect to a background model (Figure S1a). This is important, as novel driver missense mutations do not need to receive as high of a score as well-known driver missense mutations, such as KRAS G12D mutations in pancreatic ductal adenocarcinoma(Biankin et al., 2012) or IDH1 R132H mutations in gliomas (Parsons et al., 2008), but rather just need to be significantly above background. The resulting P-value distribution from CHASMplus suggest our statistical model is well calibrated (Figure S1f).

The background model was established by a somatic mutation simulation procedure as previously reported (Tokheim et al., 2016b). Briefly, using a sequence-context specific model, we repeated simulations in each gene 10 times. For each simulation, all features were computed (probabilistic2020 python package, v1.2.0). Next, each simulated missense mutation and gene was scored with the CHASMplus and 20/20+ models that were previously trained on the observed data. The resulting CHASMplus and gwCHASMplus scores for all simulations were used as an empirical null distribution. To compute a P value for a score, we used the fraction of simulated mutations with a score equal to or greater than the actual score. P values were adjusted by the Benjamini–Hochberg method for multiple hypotheses. We considered a missense mutation to be significant at a q-value threshold of 0.01.

### Compared methods

We compared CHASMplus to 12 other methods that were designed to prioritize likely cancer driver missense mutations or have been used for that purpose (VEST(Carter et al., 2013), CADD(Kircher et al., 2014), FATHMM cancer(Shihab et al., 2013), SIFT(Ng and Henikoff, 2001), MutationAssessor(Reva et al., 2011), REVEL(Ioannidis et al., 2016), MCAP(Jagadeesh et al., 2016), ParsSNP(Kumar et al., 2016), CHASM(Carter et al., 2009), Polyphen2(Adzhubei et al., 2010), transFIC(Gonzalez-Perez et al., 2012) and CanDrA(Mao et al., 2013)). We used ANNOVAR to obtain scores for 7 of the methods (VEST, CADD, SIFT, MutationAssessor, REVEL, MCAP, and Polyphen2) from dbNSFP using the ljb26_all annotation, except for REVEL and MCAP, which we used the revel and mcap annotations, respectively. TransFIC was obtained (http://bbglab.irbbarcelona.org/transfic/home) and run locally, using the scores from SIFT as input. In benchmarks specific to a cancer type, the relevant cancer type-specific model was used for CanDrA v1.0. For pan-cancer predictions, two versions of CanDrA were tested (version 1.0 and version plus, http://bioinformatics.mdanderson.org/main/CanDrA). With version plus, we used the “cancer-in-general” scores, but this was not available for version 1.0, so instead we used the ovarian scores, as these performed best. CHASM was run using a 10-fold gene-holdout cross-validation procedure using the relevant cancer type-specific model or, for pan-cancer predictions, the ovarian model. We executed ParsSNP using the provided precomputed model. FATHMM cancer scores were obtained directly from (http://fathmm.biocompute.org.uk/). Inputs to each of the methods were prepared using custom python scripts.

### Driver mutation benchmarks

We benchmarked CHASMplus in both a cancer type-specific and pan-cancer manner. In each benchmark, we defined a ‘positive’ (more driver-like) and ‘negative’ (more passenger-like) class for mutations to evaluate performance by the area under the Receiver Operating Characteristics curve (auROC). The annotation of class and mutation data used for each benchmark is listed below. Only missense mutations were used for each of the benchmarks. For reproducibility, all data, results, and analysis code are available on github (https://github.com/KarchinLab/Tokheim_2018).

#### Cancer type-specific benchmarks

We selected cancer type-specific benchmarks to evaluate how well methods could identify driver mutations relevant for specific cancer types as opposed to others. Benchmarks which contained mutations only from a single cancer type or in a single gene that is a pervasive pancancer driver gene, such as *TP53*, were not used. We chose two benchmarks based on data independent of the TCGA: cell viability of MCF10A cells and oncogenic mutations annotated by OncoKB on the MSK-IMPACT gene panel, described below. Mutations were scored using the corresponding cancer type models of CHASMplus, CanDrA, and CHASM, along with two high-performing methods which are not cancer type-specific (ParsSNP and REVEL).

##### Cell viability of MCF10A cells

MCF10A cells, a breast epithelium cell line, were used to assess cell viability of 698 missense mutations by Gordon Mills and colleagues (Ng et al., 2018). To assess each method’s ability to distinguish breast cancer-specific driver mutations, mutations that increased cell viability in known breast cancer driver genes were labeled as positive class. Mutations that did not increase cell viability or increased cell viability but were not found in breast cancer-specific genes were labeled as negative class. The latter likely represent pan-cancer drivers.

Breast cancer-specific genes were labeled based on the Cancer Gene Census (CGC, genes marked as relevant to “breast” cancer and somatic missense mutations, COSMIC v79) (Forbes et al., 2017) or The Cancer Genome Atlas (TCGA) (Network, 2012b). While Mills and colleagues (Ng et al., 2018) also evaluated a pro-B cell (Ba/F3), an appropriate cancer type-specific model was not available for all methods.

##### MSK-IMPACT gene panel mutations and OncoKB

We obtained all missense mutations from targeted sequencing with the MSK-IMPACT gene panel of 414 cancer-related genes originating from approximately 10,000 patients’ tumors (Zehir et al., 2017). Mutations annotated as ‘Oncogenic’ or ‘Likely Oncogenic’ in OncoKB (downloaded 4/3/2017) (Chakravarty et al., 2017) were selected (n=1,194). We chose four cancer types for which CanDrA v1.0 and CHASM had cancer type-specific models and which were sequenced by TGCA: Breast Invasive Ductal Carcinoma (BRCA), Glioblastoma Multiforme (GBM), Colon Adenocarcinoma (COAD), and High-Grade Serous Ovarian Cancer (OV). For each cancer type, mutations were labeled as ‘positive’ class if they occurred in a cancer driver gene implicated for that cancer type and also a tumor of the same type. The remaining mutations were then labeled as ‘negative’ class. Cancer-specific driver genes were labeled based on evidence from the CGC or TCGA. Specifically, for the CGC, we required that a gene be marked as “breast” for BRCA, as “glioma” or “glioblastoma” for GBM, as “colon” or “colorectal” for COAD, and as “ovarian” for OV, except for the exclusion of a different subtype of clear cell ovarian. Cancer-specific driver genes from the TCGA were defined based on the published marker papers for their respective cancer types: BRCA (Network, 2012b), GBM (Brennan et al., 2013), COAD (Network, 2012a), and OV (Network, 2011).

#### Pan-cancer benchmarks

We selected 5 benchmarks to evaluate the performance of CHASMplus as a pan-cancer driver predictor. CHASMplus had the highest area under the Receiver Operating Characteristics curve (auROC) of the 12 compared methods and on each of the pan-cancer benchmarks (p<0.05, DeLong test, Table S2, Figure S2b-f).

##### CGC-recurrent

We examined TCGA driver mutation prioritization at exome-wide scale through a combined literature/heuristic evaluation. We first obtained a set of curated likely driver genes from the Cancer Gene Census (CGC, COSMIC v79) (Forbes et al., 2017). Only CGC genes that were labeled as somatic and marked as relevant for missense mutations were included. We labeled all recurrent missense mutations (n>1) in the CGC genes as the positive class, and remaining mutations as the negative class (Forbes et al., 2017).

##### MSK-IMPACT gene panel mutations and OncoKB

We obtained all missense mutations from targeted sequencing with the MSK-IMPACT gene panel of 414 cancer-related genes, originating from approximately 10,000 patients’ tumors (Zehir et al., 2017). Mutations annotated as ‘Oncogenic’ or ‘Likely Oncogenic’ in OncoKB (downloaded 4/3/2017) (Chakravarty et al., 2017) were labeled as the positive class and all other mutations were labeled as the negative class.

##### Pooled *in vivo* screen in mice

A previous study by Kim et al (Kim et al., 2016) used a competitive screen of mutations in mice to assess the oncogenicity of mutations. The study selected mutations based on their presence in sequenced human tumors. Mutations were then transduced into HA1E-M cells, and pools of cells with different mutations were injected into mice and later assessed for the abundance of the mutation. Seventy-one promising alleles were then subsequently validated, individually, from the screen in NCR-Nu mice. We directly used the annotation of ‘functional’ (positive class) and ‘neutral’ (negative class) from the authors.

##### TP53 transactivation from the IARC TP53 database

We assessed each method’s ability to distinguish *TP53* mutations with low transactivation (positive class) versus all other *TP53* mutations (negative class). We evaluated all missense mutations (n=2,314) for *TP53* from the IARC *TP53* database (Petitjean et al., 2007). Low transactivation was considered as less than 50% wildtype, as indicated by the median of 8 different targets (WAF1, MDM2, BAX, h1433s, AIP1, GADD45, NOXA, and P53R2).

##### Cell viability *in vitro* assay

We evaluated missense mutations (n=747) from a medium-throughput *in vitro* experiment on two growth-factor dependent cell lines, Ba/F3 and MCF10A(Ng et al., 2018). We assessed each method’s ability to distinguish mutations resulting in increased cell viability (labeled ‘activating’; positive class) versus those that did not (labeled ‘neutral’; negative class). The experiment assumes that mutations that provide a growth advantage to cells with growth factors withdrawn reflect cancer drivers. The study considered mutations as validated if the cell viability with the mutation was higher than wild type in either cell line (2 negative controls, 3 positive controls, and wild type).

### Pan-cancer performance analysis based on area under the Precision-Recall curve

An alternative performance metric to the area under the Receiver Operating Characteristic curve (auROC) for a binary classification task is the area under the Precision-Recall curve (auPR). Like auROC, auPR summarizes the performance over all possible score thresholds from a method (Davis and Goadrich, 2006). However, auPR is preferable when there is substantial class imbalance, *i.e*., when the positive class of interest (in our case, cancer drivers) is the substantial minority (Saito and Rehmsmeier, 2015). The maximum auPR is 1.0 but the baseline score for a random predictor changes depending the skew of the class distribution (Saito and Rehmsmeier, 2015). Specifically, the expected auPR performance of a random baseline predictor is as follows:

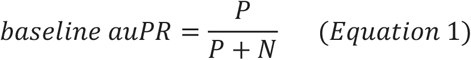

where P is the number of samples from the positive class of interest and N is the number from the other, negative class. In contrast to the auROC, auPR will give systematically high values for all methods in benchmarks that contain a majority of positive class examples (Eq. 1). Consequently, Precision-Recall curves and auPR are a poor means of comparison for several benchmarks that we used, with auROC being a better alternative.

We therefore performed auPR analysis on the two benchmarks with a substantially under represented positive class: MSK-impact gene panel (OncoKB) and CGC-recurrent (see methods). Like auROC, the auPR from the MSK-impact gene panel benchmark indicated CHASMplus had higher performance than other methods (Figure S2i). At first glance, the auPR on the CGC-recurrent benchmark for FATHMM seemed to be higher than for CHASMplus (Figure S2g). However, the performance of FATHMM and CanDrA plus dropped substantially if TP53 mutations were not included in the benchmarks (Figure S2h). This suggests the two methods may have overfit to TP53 and do not generalize as well to other genes. CHASMplus, on other hand, maintains a high auPR when TP53 is excluded, which is twice as high as the next best method.

### Gene ontology enrichment

We performed gene ontology analysis on the set of 75 genes (Table S4) containing at least one driver missense mutation from CHASMplus that did not overlap with previous genes from the Cancer Gene Census (COSMIC v79, annotated with somatic missense) or any driver genes from the TCGA PancanAtlas analysis (Bailey et al., 2018). The 75 genes were submitted to the DAVID v6.8 webserver (https://david.ncifcrf.gov/home.jsp) using default parameters for gene enrichment analysis of the biological processes gene ontology (Huang et al., 2008).

### Calculation of driver mutation frequency

Mutation frequency was calculated based on the fraction of cancer samples that contained a driver mutation in a particular cancer type. Estimates for pan-cancer analysis (32 cancer types) were based on the maximum frequency observed over the cancer types individually. The mutation frequency calculation uses the sum of driver mutations observed within the same codon. All mutations within a codon are then classified as rare (<1% of cancer samples), intermediate (1-5%), or common (>5%). Singleton mutations were considered rare, regardless of their frequency.

### CASP8 mutations and immune-related biomarkers

All values for leukocyte fraction, type of immune response (CD8 T cell, regulatory T cell, Th1 response, Th2 response, and Th17 response), and immune-related gene expression for tumor samples were obtained from (Thorsson et al., 2018). Leukocyte fraction is an estimated proportion of cells in the tumor sample that are leukocytes, as inferred from DNA methylation (Thorsson et al., 2018). The fraction consisting of CD8 T cells or regulatory T cells was estimated using the method CIBERSORT (Newman et al., 2015). Th1, Th2, and Th17 response scores are computed from RNA-Seq gene expression using single sample Gene Set Enrichment Analysis (ssGSEA) (Hanzelmann et al., 2013). Gene expression for *CTLA4, CD8A, PDCD1, CD274, TRIM1*, and *CD68* are quantitated from RNA-Seq using RSEM (score version 2)(Li and Dewey, 2011).

### Comparison with PTEN multiplexed functional assays

CHASMplus was compared to two previous multiplexed functional assays of PTEN, examining lipid phosphatase activity (Mighell et al., 2018) and intracellular protein abundance (Matreyek et al., 2018). We first compiled a list of 6,564 missense mutations designated as having high confidence measurements of lipid phosphatase activity by Mighell and colleagues. We found the overlap with those measured by Matreyek *et al*., resulting in 3,540 missense mutations with both lipid phosphatase activity and intracellular protein abundance. gwCHASMplus scores were computed for each of these missense mutations. Correlation of gwCHASMplus with functional scores of lipid phosphatase activity and protein abundance was computed using Locally Weighted Scatterplot Smoothing (LOWESS) and spearman rank correlation. Driver missense mutations identified by (pan-cancer) CHASMplus were compared to all other missense mutations observed in PTEN using a two-sided Mann-Whitney U test. Finally, we calculated auROC (Figure S6) to assess whether the functional scores could discriminate between ClinVar labeled pathogenic or likely pathogenic variants and variants found in gnomAD (n=163).

We extended our comparison of the PTEN saturation mutagenesis experiment of lipid phosphatase activity to the two “runner-up” methods (CanDrA and ParsSNP), assessed in the five benchmarks in Figure 2d. Mutations were regarded as damaging to lipid phosphatase activity if the functional score from the multiplexed functional assay was below the recommended value of −1.11 (Mighell et al., 2018). To dichotomize the predictions of CanDrA and ParsSNP, we used the driver calls from CanDrA and the recommended threshold value of 0.1 by ParsSNP. By necessity, analysis was restricted to the set of missense mutations that are formed by single base substitutions (n=2,079), as these are the only missense mutations that can be handled by CanDrA and ParsSNP. CHASMplus, CanDrA and ParsSNP were then compared to each other based on their specificity, sensitivity, precision and F1 score to identify mutations damaging to lipid phosphatase activity in PTEN.

### Clustering of cancer types

We clustered TCGA cancer types according to two features, prevalence (fraction of samples mutated) and normalized diversity (normalized entropy) among predicted missense mutation drivers (q <= 0.01). The normalized entropy score was calculated based on the codon-level, as follows,

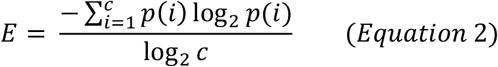

where there are c codons containing significant mutations, and the fraction of significant mutations in the i’th codon is p(i). We performed clustering using the k-means algorithm (scikit learn v0.18.0) where k, the number of clusters, was selected by the maximum silhouette score (k=5; varied between 2 and 10). Each parameterization was run ten times with different initial conditions to avoid local optimums by choosing the best run, defined as the lowest sum of distances to the closest centroid.

Beyond biological differences intrinsic to each cancer type, technical difficulties in mutation calling could possibly explain the above clustering patterns. To evaluate this possibility, we correlated the mean Variant Allele Fraction (VAF) for mutations in tumor samples in each cancer type with a variety of metrics summarizing our results. VAF acts as a combined indicator of mutation sub-clonality and normal tissue contamination within the tumor sample (Cibulskis et al., 2013), both of which lower the capability to detect mutations. We found no significant correlation between mean VAF for cancer types with: average number of predicted driver mutations per sample (Pearson r=0.26, p=0.14, correlation test), fraction of samples with predicted driver mutations (Pearson r=0.2, p=0.28, correlation test), unique number of significant mutations (Pearson r=0.04, p=0.82, correlation test), and normalized driver diversity (Pearson r=0.33, p=0.07, correlation test).

### Subsampling procedure

We performed driver missense mutation predictions on random subsamples of each of 9 representative cancer types (ACC, SARC, PRAD, THYM, UVM, PAAD, BRCA, HNSC, and COAD), using CHASMplus. Subsampling was performed by randomly selecting a certain fraction of tumor samples without replacement. The designated fraction of samples for each iteration was randomly selected from a uniform distribution bounded between 0 and 1. We then ran CHASMplus using a 10-fold gene-holdout cross-validation model previously trained on the TCGA mutation data. The number of unique driver missense mutations and overall driver prevalence (average number of driver missense mutations per cancer sample) were then calculated based on significant CHASMplus predictions (q<=0.01). The prevalence within a particular sub-sampled result was measured against the full cohort. Results were then ordered by increasing sample size to observe trends in the identification of driver missense mutations by CHASMplus.

### Analysis of the tail of driver discovery for Prostate Adenocarcinoma

We used mutations from a meta-analysis of 1,013 prostate adenocarcinoma (PRAD) samples (Armenia et al., 2018), which included TCGA and several other studies, to assess our predicted trajectory of discovery of driver missense mutations in PRAD from TCGA (n=477), Genes that contained putative driver missense mutations predicted by CHASMplus substantially overlaped the significantly mutated genes identified by the authors of this study (Figure S7b). They reported a larger number of genes, but this included genes driven by mutations other than missense mutations (e.g., nonsense, frameshift, splice site, etc.), genes from other types of cancers, and a much higher FDR (25% vs 1% for CHASMplus). Based only on TCGA, we predicted a sub-linear increase in prevalence of driver missense mutations in PRAD (Figure 6b). After applying FDR correction at 1%, driver prevalence in the larger study was 0.4 mutations per sample compared to 0.31 mutations per sample in the TCGA PRAD cohort. A strictly linear increase from 477 to 1,013 samples would have yielded 0.66 mutations per sample, supporting our prediction of a sub-linear increase.

### Limited power for mutation hotspot detection approaches

A codon or small region of protein sequence or structure where recurrent mutations are observed is known as a hotspot. Similar to statistical methods for driver gene detection, hotspot detection identifies an excess number of mutations compared to expectation using a large number of cancer samples. We asked whether, given current cohort sizes, codon-based hotspot detection had sufficient statistical power to identify rare driver mutations. We assessed the number of samples required to detect driver missense mutations across a range of frequencies (proportion of tumor samples in which a mutation occurs) and somatic background mutation rates. In Figure 7a, each of the 32 TCGA cancer types is placed according to its sample size and background mutation rate, relative to six curves which represent the required sample size to detect driver missense mutations of a certain frequency, with 90% power, using hotspot detection (see Statistical Power Analysis). For example, the TCGA Cervical Squamous Cell Carcinoma and Endocervical Adenocarcinoma (CESC) cohort has 274 samples and a background mutation rate of 3.5 mutations/Mb. This sample size is sufficient to detect driver missense mutations that occur in ~2% of the samples with 90% power. At current TCGA sample sizes, we found codon-based hotspot detection approaches were not well powered to identify driver missense mutations that occurred at less than 1% frequency in most cancer types. Exceptions were thyroid carcinoma (THCA), low grade glioma (LGG) and breast cancer (BRCA), which are seen to lie above (or close to) the curve representing 1% frequency (Figure 7a). Notably, these cohorts had large numbers of samples and low-to-medium background mutation rates. We also found that when cancer types were aggregated in pan-cancer analysis, power to detect codon-based hotspots improved substantially, but only when the recurrent mutations were shared in more than one cancer type. For these mutations, pan-cancer analysis using ~10,000 TCGA samples should enable detection of driver mutations at frequency as low as 0.1%.

### CHASMplus has greater power than a hotspot detection method

We compared (pan-cancer) CHASMplus to the cancer hotspots method (v0.6)(Chang et al., 2016), with respect to its sensitivity to identify driver missense mutations. Cancer hotspots was download from github (https://github.com/taylor-lab/hotspots) and run using default parameters on the full TCGA mutation dataset. For each gene, we used the biomart R package to compute the length of its protein product. For both methods, we computed the overlap of missense mutations called as driver or hotspot (q<=0.01) with well-curated oncogenic mutations in the OncoKB database. The sensitivity of CHASMplus to detect the OncoKB-labeled mutations was 0.83, which was significantly higher than the hotspot method (0.46, p<2.2e-16, McNemar’s test, n=896). To minimize potential gene bias, we also repeated the analysis after excluding all 389 TP53 mutations, yielding sensitivity of 0.76 for CHASMplus and 0.49 for hotspot detection, a difference which is still statistically significant (p<2.2e-16, McNemar’s test, n=507) (Figure 7b). The increased sensitivity did not come at the cost of low specificity, as evidenced by our p-value calibration (Figure S1f) and extensive ROC analysis across five benchmarked datasets (Figure 2), which measures a balance of sensitivity and specificity.

### Calibration of p-values

Most driver missense mutation prediction methods do not report p-values, but CHASMplus and cancer hotspots (Chang et al., 2016) do compute a p-value. P-values under the null hypothesis (in this case, passenger events) are expected to follow a uniform distribution (Storey and Tibshirani, 2003). We therefore compared the distribution of p-values produced by these methods to a uniform distribution by using a QQ plot (Wilk and Gnanadesikan, 1968), which compares the quantiles of two distributions. We removed likely driver events by excluding all p-values from genes included in the Cancer Gene Census, a well-curated collection of driver genes (Futreal et al., 2004). A well-calibrated statistical model should closely follow the diagonal on a QQ plot, given that driver events in tumors are few compared to passengers (Tomasetti et al., 2015; Vogelstein et al., 2013), especially when mutations in known cancer driver genes are removed.

### Statistical power methodology

We estimated the required statistical power to find codons with mutation frequency significantly above background. We used a binomial model previously developed for driver gene power analysis (Figure 7a) (Lawrence et al., 2014; Tokheim et al., 2016b). A gene-specific mutation rate factor *F_g_* was set to represent a gene at the 90th percentile, given an exome-wide background mutation rate of *π*, so that *μ* = *F_g_π* (*F_g_* = 3.9). Because our analysis was based on codons, the length parameter L was set to that of a codon *L_c_ =* 3. We assumed that ~¾ of mutations to be non-silent as in (Lawrence et al., 2014), so the effective gene length was adjusted as 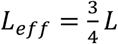. Background mutation rate was calculated using the total number of potentially mutated bases that could yield a non-silent mutation (*N_eff_*), which is the effective length multiplied by number of samples (S). For each codon of interest, we compared the null hypothesis to the alternative hypothesis that its mutation frequency exceeds the background. A Bonferroni correction was applied at a family-wise error rate of 0.1. A codon with significantly higher non-silent mutation rate per-base (*μ_es_*) than that codon’s background mutation rate *μ* is defined as

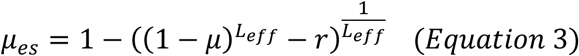

and *r* is the fraction of samples with non-silent mutations in the codon above background. Samples were iteratively added until there was greater than or equal to 90% probability that a driver gene with mutation rate *μ_es_* would be found significant.

